# Arabidopsis poly(ADP-ribose)-binding protein RCD1 interacts with Photoregulatory Protein Kinases in nuclear bodies

**DOI:** 10.1101/2020.07.02.184937

**Authors:** Julia P. Vainonen, Alexey Shapiguzov, Julia Krasensky-Wrzaczek, Richard Gossens, Raffaella De Masi, Iulia Danciu, Tuomas Puukko, Natalia Battchikova, Claudia Jonak, Lennart Wirthmueller, Michael Wrzaczek, Jaakko Kangasjärvi

**Author notes:** Corresponding author: Jaakko Kangasjärvi. These authors contributed equally to the manuscript (Julia P. Vainonen, Alexey Shapiguzov, Julia Krasensky-Wrzaczek, Richard Gossens).

## Abstract

Continuous reprogramming of gene expression in response to environmental signals is required for plant survival in changing environment. One mechanism responsible for this is signaling through hub proteins that integrate external stimuli and transcriptional responses. RADICAL-INDUCED CELL DEATH1 (RCD1) functions as a nuclear hub protein, that interacts with a variety of transcription factors through its C-terminal RST domain and acts as a co-regulator of numerous stress responses in plants. Here, a previously unknown function for RCD1 as a novel plant poly(ADP-ribose) (PAR) reader protein is described. RCD1 localizes to specific locations inside the nucleus, in a PAR-dependent manner; its N-terminal WWE domain o binds PAR and together with the PARP-like domain determines its localization to nuclear bodies (NBs), which is prevented by inhibition of PAR synthesis. RCD1 also interacts with Photoregulatory Protein Kinases (PPKs) that co-localize with RCD1 in the NBs. The PPKs, that have been associated with circadian clock, abscisic acid, and light signaling pathways, phosphorylate RCD1 at multiple sites in the intrinsically disordered region between the WWE and PARP-like domains. This affects its stability and functions in the nucleus and1 provides a mechanism where the turnover of a PAR-binding transcriptional co-regulator is controlled by nuclear protein kinases.

## INTRODUCTION

Plants are constantly exposed to a variety of environmental cues that are relayed to the nucleus, which coordinates adaptation to challenges associated with those cues. The eukaryotic nucleus has many functions including DNA and RNA biogenesis and processing, transcriptional regulation, RNA splicing, protein modification and degradation. Many functions are organized within non-membranous compartments, so-called “nuclear bodies” (NBs; Mao *et al*, 2011). Several types of NBs have been identified, such as the nucleolus, Cajal bodies, Polycomb bodies, and photobodies. Specific NB-associated proteins have also been described, including members of the splicing machinery (Reddy *et al*, 2012), chromatin-associated proteins (Simon *et al*, 2015), ubiquitin ligases (Christians *et al*, 2012), photoreceptors (Van Buskirk *et al*, 2012), and protein kinases (Wang *et al*, 2015).

*Arabidopsis thaliana* RADICAL-INDUCED CELL DEATH1 (RCD1) is a nuclear-localized multidomain protein comprised of an N-terminal bipartite nuclear localization sequence (NLS), a WWE domain, a poly(ADP-ribose) polymerase-like (PARP-like) domain, and an RCD1-SRO-TAF4 (RST) domain (Ahlfors *et al*, 2004; Jaspers *et al*, 2009; Jaspers *et al*, 2010; **Figure 1A**). The domains of RCD1 are flanked by intrinsically disordered regions (IDRs), which likely provide flexibility in assuming the final overall protein conformation. Arabidopsis RCD1 and its paralog, SRO1 (SIMILAR TO RCD ONE 1), can form homo- and heterodimers (Wirthmueller *et al,* 2018). The presence of at least one of the paralogs is essential since the *rcd1 sro1* double mutant displays defects in embryo development and is not viable under standard growth conditions (Jaspers *et al*, 2009; Teotia & Lamb, 2011).

**Figure 1.**
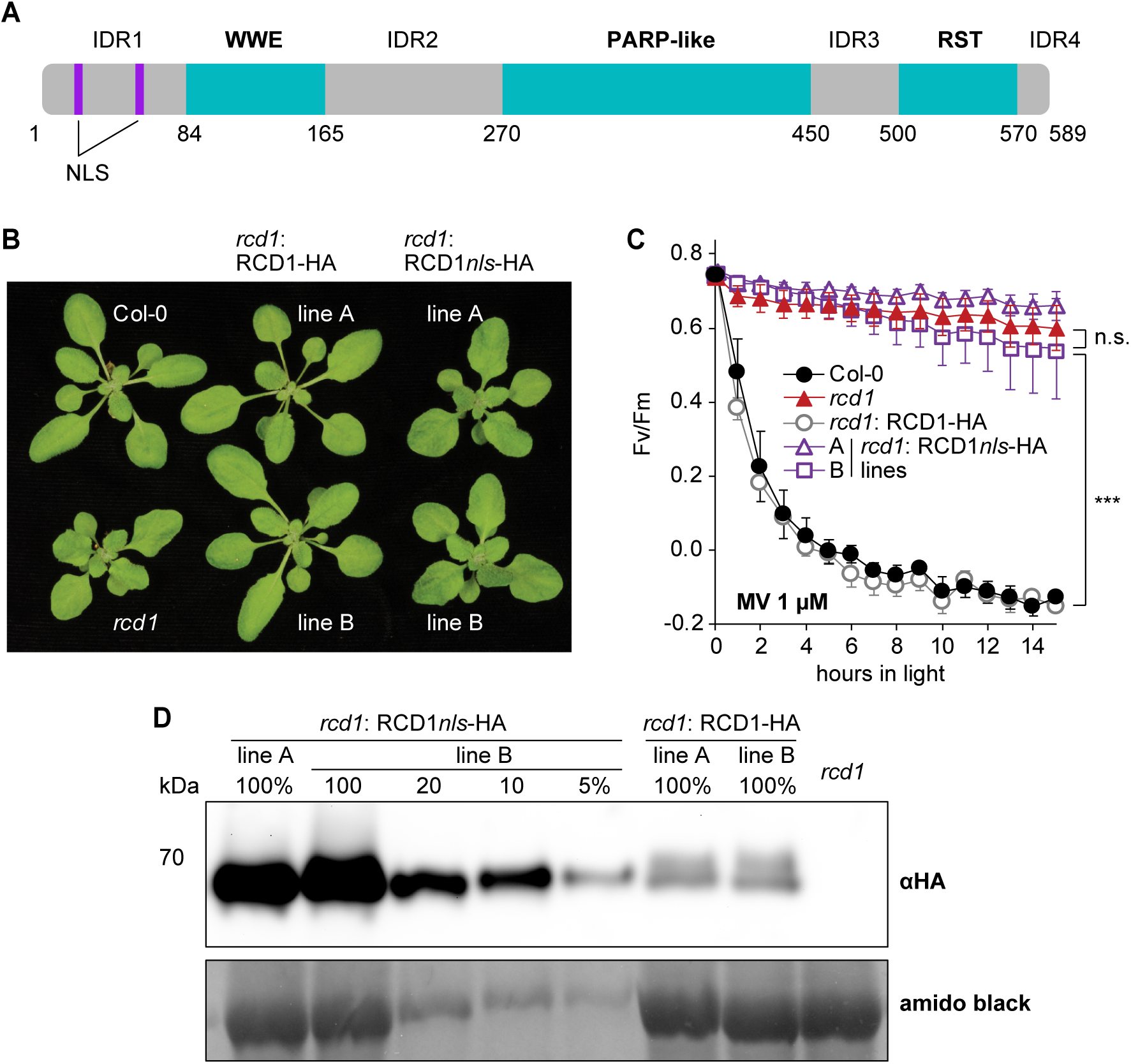
Nuclear localization of RCD1 is essential for its function. **A.** Schematic representation of RCD1 domain structure containing a bipartite NLS, WWE, PARP-like and RST domains. Intrinsically disordered regions between the domains are marked as IDR1-4. **B.** Curly leaf phenotype of *rcd1* can be complemented by re-introduction of wild type RCD1-HA, but not of RCD1 with mutated NLS (RCD1*nls*-HA) into the mutant background. The photo shows 3-week-old plant rosettes of two independent lines (A and B) for each construct under standard growth conditions. **C.** RCD1 requires its NLS to complement the *rcd1*-specific MV tolerance. PSII inhibition (Fv/Fm) by methyl viologen (MV) was measured in indicated lines using 1 μM MV. For each experiment, leaf discs from three individual rosettes were used. The experiment was performed three times with similar results. Mean ± SD are shown. *** – P-value < 0.001 with Tukey corrected *post hoc* test; n.s. – non-significant difference. Source data and statistics are presented in **Supplementary table 1**. **D.** Disruption of NLS leads to higher RCD1 accumulation in plants. Abundance of RCD1-HA in RCD1*nls*-HA and RCD1-HA lines was assessed by immunoblot analysis with HA-specific antibodies. A total protein amount of 100 µg corresponds to 100%. Rubisco large subunit detected by amido black staining is shown as a control for protein loading.

In plants, the RST-domain is unique to the RCD1-SRO protein family and TAF4 proteins (Jaspers *et al*, 2010). It has been described as a domain that mediates interactions with many RCD1-associated proteins (O’Shea *et al*, 2017; Bugge *et al*, 2018; Shapiguzov *et al*, 2019). A structurally diverse set of transcription factors interacts with the RST domain, making RCD1 an important hub for transcriptional regulation (Jaspers *et al*, 2009; Vainonen *et al*, 2012; O’Shea *et al*, 2017; Christensen *et al*, 2019; Shapiguzov *et al*, 2019; Jespersen & Barbar, 2020). Unlike the RST domain, the WWE and PARP-like domains of RCD1 have hardly been characterized.

The WWE domain was originally defined computationally and proposed to be a protein-protein interaction domain in proteins related to ubiquitination and ADP-ribosylation (Aravind 2001). Later studies have shown that some, but not all animal WWE domains bind iso-ADP ribose, a structural unit of poly(ADP-ribose) – PAR (Wang *et al*, 2012; DaRosa *et al*, 2015). In *Arabidopsis thaliana*, the WWE domain has only been identified in RCD1 and SRO1. While the PARP-like domain in these proteins does not exhibit detectable PARP activity (Jaspers *et al*, 2010; Wirthmueller *et al,* 2018), the presence of WWE and a PARP-like domains together suggests a function of RCD1 in PAR-related processes (Vainonen *et al*, 2016).

Poly-ADP-ribosylation (PARylation) of proteins is a reversible posttranslational modification, which has been intensively studied in animals during recent decades (Gupte *et al*, 2017; Cohen & Chang, 2018). PARPs catalyze PARylation by covalently attaching ADP-ribose moieties to glutamate, aspartate, lysine, arginine, serine, threonine and cysteine residues in a species- and tissue-specific manner (Jungmichel *et al*, 2013; Zhang *et al*, 2013; Martello *et al*, 2016; Leung 2017; Palazzo *et al*, 2018). PAR-glycohydrolase (PARG) can trim down PAR chains to the terminal protein-bound ADP-ribose thereby removing PAR from proteins. Additionally, several signaling components that recognize PARylated proteins, so-called “PAR readers”, have been identified in animal systems (Gupte *et al*, 2017; Kim *et al*, 2020), but have not been described in plants yet. On a functional level, in animal cells PARylation has been shown to regulate a variety of cellular processes including chromatin remodeling, transcription, and programmed cell death (Gupte *et al*, 2017; Kim *et al*, 2020). In plants, the role of PAR is only starting to emerge and the few studies available suggest an important role for PARylation in plant stress and developmental responses (Vainonen *et al*, 2016).

In addition to transcription factors, RCD1 has been shown to interact with Photoregulatory Protein Kinases (PPKs; also named MUT9-like kinases, MLKs, or Arabidopsis EL1-like kinases, AELs; Wirthmueller *et al,* 2018; Shapiguzov *et al,* 2019). In Arabidopsis, this recently discovered family of protein kinases is comprised of four members that localize to NBs (Wang *et al*, 2015). PPKs interact with different nuclear proteins, including histones, components of the circadian clock and light signaling, and the ABA receptor PYR/PYL/RCAR (Wang *et al*, 2015; Huang *et al*, 2016; Liu *et al*, 2017; Ni *et al*, 2017; Su *et al*, 2017; Chen *et al*, 2018; Zheng *et al*, 2018). PPK-dependent phosphorylation of the transcription regulators PIF3 and CRY2, and the ABA receptor PYR/PYL/RCAR has been shown to target these proteins for degradation (Liu *et al*, 2017; Ni *et al*, 2017; Chen *et al*, 2018). However, the mechanistic roles for phosphorylation of histone and circadian clock component by PPKs have not been described so far.

Here we show that Arabidopsis RCD1 localizes to NBs in a PAR-dependent manner. RCD1 binds PAR via the WWE domain and can be described as the first identified PAR reader in plants. Furthermore, we demonstrate that RCD1 is phosphorylated by members of the PPK protein kinase family, which co-localize with RCD1 *in vivo* in NBs. Increased RCD1 protein levels together with altered tolerance to oxidative stress in *ppk* mutant plants suggest that phosphorylation by PPKs regulates RCD1 protein stability.

## RESULTS

### Nuclear localization is essential for RCD1 function

To address the role of subcellular localization for the function of RCD1, the basic amino acids of the NLS were substituted with aliphatic ones (K21L/R22I and R56I/R57I). These point mutations were introduced into a construct expressing RCD1 tagged with triple HA epitope at the protein C-terminus, under the native RCD1 promoter (Jaspers *et al*, 2009) and transformed into *rcd1* background (RCD1*nls*-HA lines hereafter). To verify that disruption of the NLS prevents nuclear localization of RCD1, we also generated stable Arabidopsis lines in *rcd1* background expressing wild type RCD1 or RCD1*nls* fused to a triple Venus tag under the control of the UBIQUITIN10 promoter (*rcd1:*RCD1-Venus and *rcd1:*RCD1*nls*-Venus lines). The subcellular localization of the tagged proteins was studied by confocal microscopy. In contrast to the wild type protein that was localized to the nucleus, RCD1*nls*-Venus was localized outside of nuclei (**Supplementary figure 1A**).

RCD1*nls*-HA lines were further studied for their ability to complement *rcd1* phenotypes. Among several stress- and development-related phenotypes (Overmyer *et al*, 2000; Ahlfors *et al*, 2004; Jaspers *et al*, 2009; Teotia &Lamb, 2009; Hiltscher *et al*, 2014), the *rcd1* mutant displays curly leaves and tolerance to the herbicide methyl viologen (MV) (Ahlfors *et al*, 2004; Fujibe *et al*, 2004; Shapiguzov *et al*, 2019). At the molecular level, MV interferes with the electron transfer chain in chloroplasts and catalyzes production of reactive oxygen species (ROS). Introduction of RCD1-HA, but not RCD1*nls*-HA, into *rcd1* restored the wild type shape of the leaves (**Figure 1B**). Similarly, in *rcd1:*RCD1*nls*-HA plants the increased tolerance of *rcd1* to MV was not reverted to wild type-like sensitivity (**Figure 1C**). Analysis of protein levels in the transgenic lines revealed increased RCD1 accumulation in RCD1*nls*-HA lines compared to wild type RCD1-HA (**Figure 1D**). Thus, despite higher levels of RCD1 in RCD1*nls*-HA lines, *rcd1* phenotypes were not complemented. These results suggest that nuclear localization is required for the proper function of RCD1.

### WWE and PARP-like domains are required for RCD1 function

The role of individual domains of RCD1 was addressed by generation of RCD1 domain deletion constructs tagged with triple HA epitope and expressed under the native RCD1 promoter (*rcd1:*RCD1ΔWWE-HA, *rcd1:*RCD1ΔPARP-like-HA, *rcd1:*RCD1ΔRST-HA lines hereafter). The structure of the domain deletion constructs is shown in **Supplementary figure 1B**. These lines were tested for characteristic *rcd1* phenotypes, including plant habitus, flowering time and MV tolerance. At least two individual lines for each protein variant were used in all the experiments.

Expression of RCD1 lacking the individual domains in *rcd1* background did not fully complement the curly leaf phenotype and early flowering specific to the *rcd1* mutant (**Figure 2A, 2D**). The abundance of RCD1-HA in the domain deletion lines was verified by immunoblot analysis with HA-specific antibody (**Figure 2B**). These analyses confirmed that the lack of complementation was not due to the absence of protein in these lines. The domain deletion lines also exhibited high abundance of the mitochondrial alternative oxidase AOX1/2 proteins **(Figure 2B**), which is characteristic for the *rcd1* mutant (Shapiguzov et al, 2019). Analysis of the lines showed that the RCD1 domain deletions did not complement the *rcd1*-specific MV tolerance to the wild type sensitivity (**Figure 2C, Supplementary figure 2**); these results are in agreement with higher AOX1/2 expression in the domain deletion lines. Deletion of the RST domain resulted in MV tolerance phenotype most similar to *rcd1*, whereas *rcd1:*RCD1ΔWWE-HA and *rcd1:*RCD1ΔPARP-like-HA lines exhibited intermediate sensitivity to MV. Only wild type RCD1-HA fully complemented the *rcd1* early flowering phenotype. RCD1ΔPARP-like-HA complemented the early flowering only partially, while RCD1ΔWWE-HA did not complement this phenotype at all. Interestingly RCD1ΔRST-HA intensified early flowering (**Figure 2D, Supplementary figure 2**).

**Figure 2.**
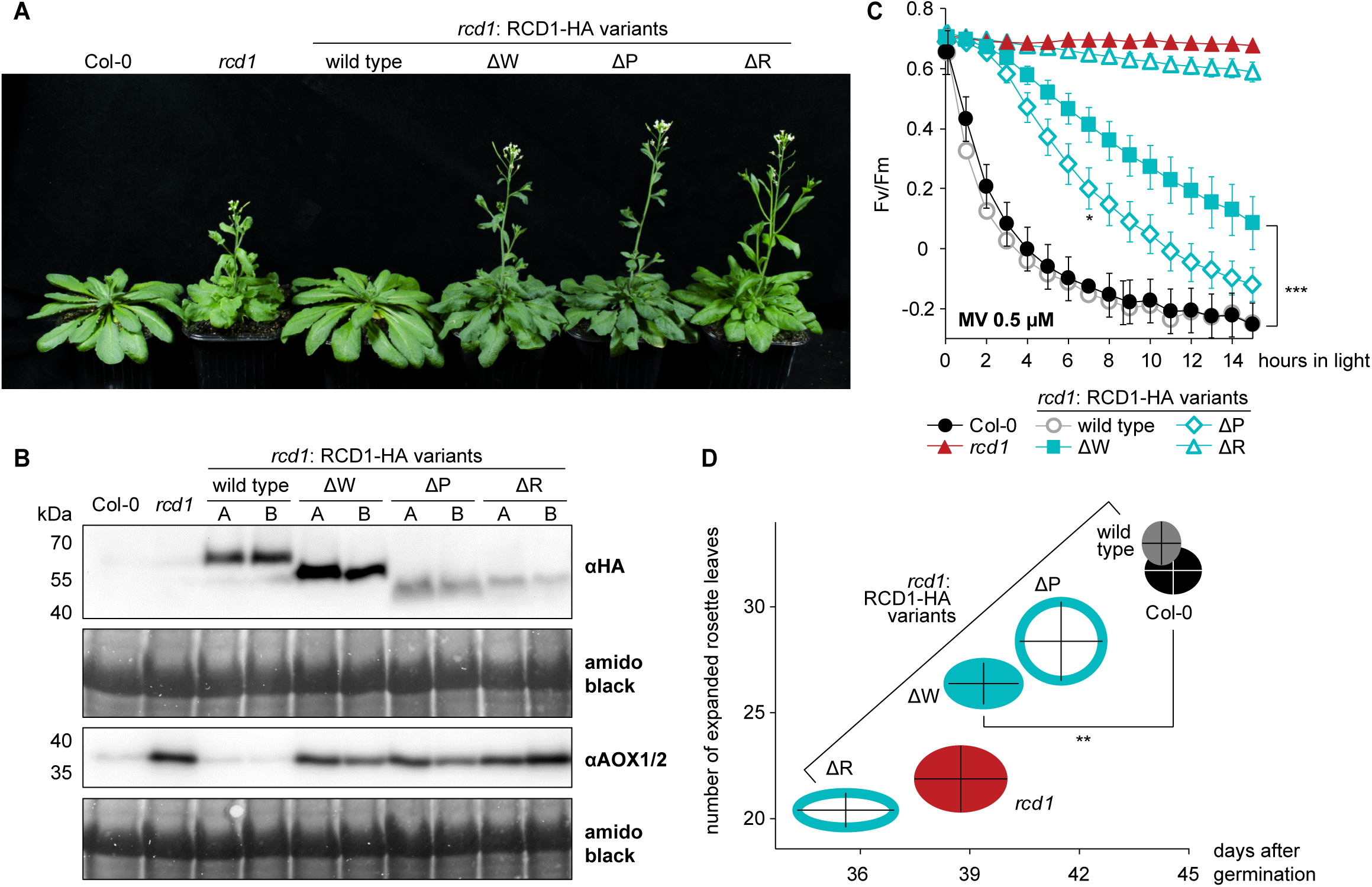
WWE, PARP-like and RST domains are necessary for RCD1 functions. **A.** Deletion of RCD1 individual domains prevents complementation of *rcd1* early flowering phenotype in Arabidopsis. Depicted are lines expressing wild type RCD1 or RCD1 versions lacking the WWE (ΔW), PARP-like (ΔP) or RST (ΔR) domains. The photo shows 5-week-old plants of representative lines under standard growth conditions. **B.** Immunoblot analysis of two independent domain deletion lines for each construct (A and B) shows presence of RCD1-HA in complementation lines (upper panel) and increased AOX1/2 expression in these lines at the level similar to the *rcd1* mutant (middle panel). Rubisco large subunit detected by amido black staining is shown as a control for equal protein loading. **C.** Wild type MV sensitivity is not restored in lines expressing RCD1ΔWWE-HA (ΔW), RCD1ΔPARP-like-HA (ΔP), and RCD1ΔRST-HA (ΔR) constructs. PSII inhibition (Fv/Fm) by MV was measured in indicated lines using 0.5 μM MV. For each experiment, leaf discs from three individual rosettes were used. The experiment was performed three times with similar results. Mean ± SD are shown. * – P-value < 0.05 with Tukey corrected *post hoc* test at the selected time point between *rcd1*: RCD1ΔPARP-like-HA and *rcd1*: RCD1-HA lines; *** – P-value < 0.001 with Tukey corrected *post hoc* test at the selected time point between *rcd1*: RCD1ΔWWE-HA and *rcd1*: RCD1-HA lines. Source data and full statistics are presented in **Supplementary table 1**. **D.** Early flowering phenotype of *rcd1* is not fully restored by RCD1-HA deletion constructs. Flowering time defined as the day of the opening of the first flower after germination, is plotted against the number of expanded rosette leaves on the flowering day. The experiment was performed three times with similar results. Mean ± SE are shown as intersection and black error bars. ** – P-value < 0.01 with Tukey corrected *post hoc* test. Source data and full statistics are presented in **Supplementary table 1**.

Together, these results indicated that the WWE and the PARP-like domains of RCD1 are required for at least some of RCD1 functions in the nucleus together with the RST domain that mediates RCD1 interaction with transcription factors.

### RCD1 localizes to NBs and binds PAR

To study the nuclear localization of RCD1 in further detail, we used Arabidopsis lines expressing wild type RCD1-Venus fusion protein described above, as well as deletion constructs lacking the individual domains (WWE, PARP-like, and RST) under control of the UBIQUITIN10 promoter in *rcd1* background (**Supplementary figure 1B**). Microscopic analysis of RCD1-Venus lines showed that RCD1 localized exclusively to the nucleus. Within the nucleus, RCD1-Venus localized to the nucleoplasm and, intriguingly, to distinct NBs (**Figure 3A**). Deletion of the WWE or the PARP-like domain, but not the RST domain, suppressed localization of RCD1 to these NBs under standard growth conditions (**Figure 3A**). Immunoblot analysis of the corresponding lines showed increased levels of RCD1-Venus in all deletion constructs compared to wild type RCD1-Venus (**Figure 3B**). Thus, the apparent inability of RCD1ΔWWE and RCD1ΔPARP-like to form NBs was not due to low protein abundance.

**Figure 3.**
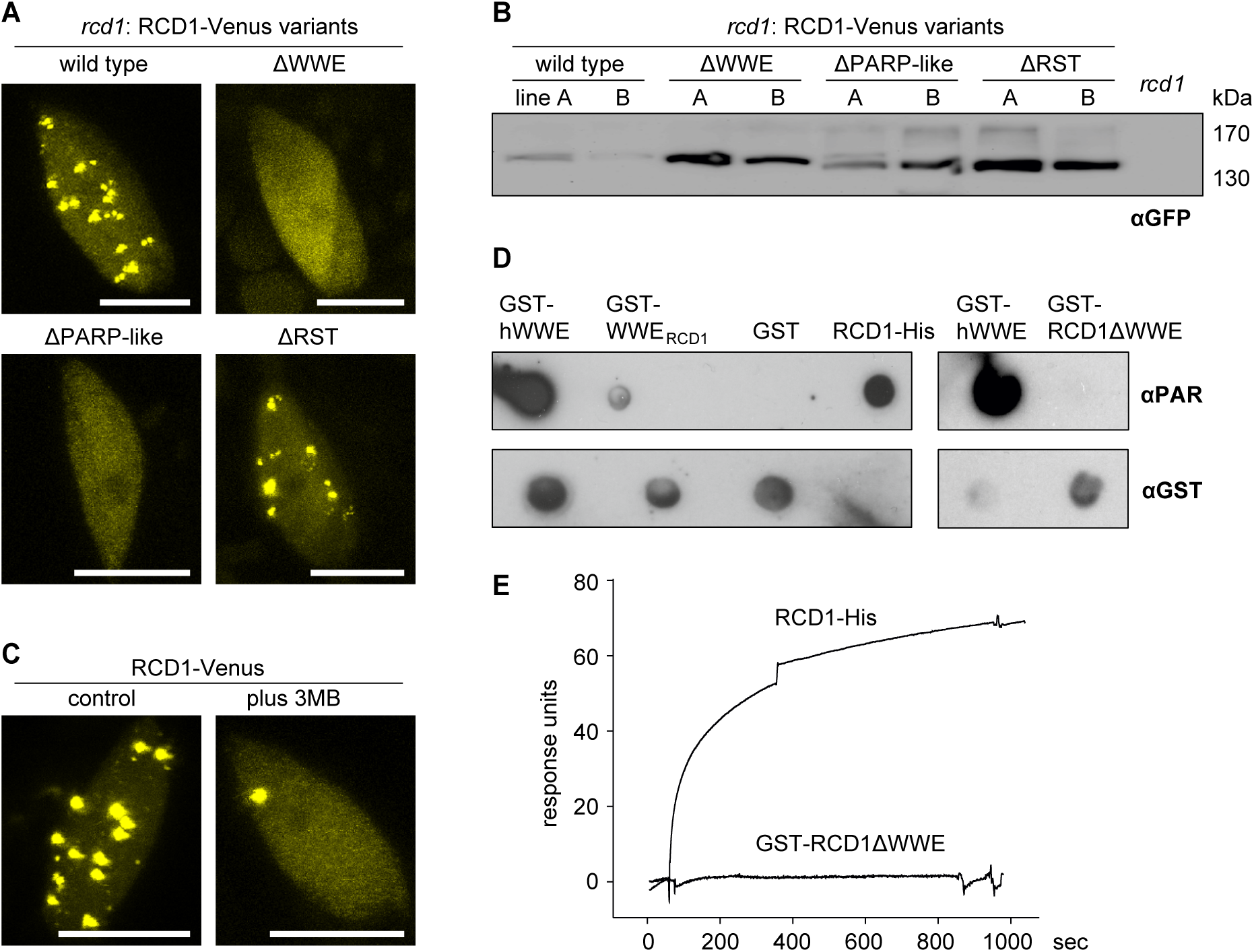
RCD1 localizes to NBs dependent on WWE and PARP-like domains and binds PAR. **A.** Deletion of the WWE or PARP-like domains, but not the RST domain, prevents NB localization of RCD1. Confocal images were taken from stable Arabidopsis lines expressing full-length RCD1-Venus, RCD1ΔWWE-Venus, RCD1ΔPARP-Venus or RCD1ΔRST-Venus in the *rcd1* background. Scale bars indicate 10 µm. **B.** Domain deletion does not lead to decreased expression of RCD1. RCD1 level in indicated lines was assessed by immunoblot analysis of total protein extracts with GFP-specific antibody. A total amount of 100 μg protein was loaded per lane. **C.** NB localization of RCD1 is diminished by PARP inhibitor 3MB. Plants expressing RCD1-Venus were pretreated overnight at 4°C without (control) or with 3MB, after which confocal microscopy was performed. Scale bars indicate 10 µm. **D.** RCD1 binds PAR *in vitro*. PAR binding activity of immobilized GST-tagged domains of RCD1 and full-length RCD1-His was assessed by dot-blot assay using PAR-specific antibody. GST tagged human WWE domain (hWWE) and GST were used as positive and negative controls, respectively. GST antibody was used to assess protein loading. **E.** WWE domain of RCD1 is required for interaction with PAR. SPR sensorgrams of interaction between immobilized RCD1-His or GST-RCD1ΔWWE and PAR profiled at 625 nM. Increase in response units shows association of PAR with RCD1-His but not with GST-RCD1ΔWWE.

The WWE domain has previously been described to bind PAR in mammalian cells (Zhang *et al*, 2011; Wang *et al*, 2012; DaRosa *et al*, 2015). Therefore, we tested whether a chemical inhibitor of PAR synthesis, 3-methoxybenzamide (3MB), would influence NB localization of RCD1-Venus. Indeed, in 3MB-treated plants, RCD1-Venus localized almost exclusively to the nucleoplasm (**Figure 3C**). This suggests that the presence of PARylated proteins in the nucleus is necessary for RCD1 to localize to NBs.

To examine whether RCD1 can bind PAR directly, we tested the interaction *in vitro*. Recombinant proteins were expressed in *E.coli*, purified (**Supplementary Figure 3A**) and dot-blotted on a nitrocellulose membrane. The membrane was then incubated with PAR polymer, washed, and subsequently probed with anti-PAR or anti-GST antibodies. As shown in **Figure 3D**, the WWE domain of RCD1 alone, as well as the full-length protein (fused to either GST or His-tags, respectively), interacted with PAR. To verify that PAR-binding was mediated by the WWE domain, we tested the PAR-binding properties of a truncated version of RCD1 lacking the WWE domain (GST-RCD1ΔWWE). This variant was not able to bind PAR as shown in **Figure 3D**. In contrast, deletion of the PARP-like domain did not prevent *in vitro* PAR binding by RCD1 in the dot-blot assay (**Supplementary figure 3B**).

For quantitative characterization of the RCD1-PAR interaction, we applied surface plasmon resonance (SPR), a method that allows label-free detection of biomolecular interactions. SPR demonstrated that full-length RCD1 interacted with PAR and that the interaction was abolished by deletion of the WWE domain (GST-RCD1ΔWWE; **Figure 3E**). The binding curve was similar with the WWE domains described in other studies (Zhang *et al*, 2011; Wang *et al*, 2012). Absence of dissociation in the running buffer (**Figure 3E**) confirmed strong complex formation between PAR and RCD1. Estimation of the affinity of the interaction by binding experiments with increasing concentrations of the PAR ligand resulted in a dissociation constant of 28.2 nM (**Supplementary Figure 3C**) indicating high-affinity interaction. Furthermore, in the SPR analyses RCD1 did not interact with compounds related to PAR, such as monomeric ADP-ribose, or cyclic ADP-ribose, a known second messenger (**Supplementary Figure 3D** and **E**). Thus, our experiments showed that RCD1 specifically binds PAR with high affinity and this interaction requires the WWE domain.

### RCD1 co-localizes with PPKs in NBs

Immunoblot analyses showed that nuclear-localized RCD1-HA migrated in SDS-PAGE as a double band (**Figure 1D**), indicative of post-translational modification of the protein. To test whether the double band was caused by phosphorylation of RCD1, protein extracts from plants expressing wild type RCD1-HA were treated with calf intestinal alkaline phosphatase. The phosphatase treatment eliminated the double band of RCD1-HA (**Supplementary Figure 4**), suggesting that RCD1 is an *in vivo* phosphoprotein. RCD1-HA migrated as a single band in the transgenic lines expressing RCD1*nls*-HA (**Figure 1D**), suggesting that nuclear localization was necessary for phosphorylation of RCD1.

Our previous proteomic analyses of RCD1 interactors (Wirthmueller *et al*, 2018; Shapiguzov *et al*, 2019), showed that RCD1 interacted *in vivo* with a newly described family of protein kinases, the Photoregulatory Protein Kinases (PPKs). This interaction was confirmed by targeted co-immunoprecipitation experiments in tobacco using RCD1-GFP and PPK-HA constructs, in which RCD1-GFP co-immunoprecipitated with PPK1, 3 and 4 (**Supplementary Figure 5**). The apparent lack of interaction between RCD1 and PPK2 in this assay could indicate either isoform-specific differences in the strength of association, or technical limitations as PPK2 protein was hardly detectable in total protein extracts. Collectively, these data confirmed complex formation between Arabidopsis RCD1 and PPKs also in a heterologous *in vivo* plant expression system.

It has previously been shown that PPKs localize to NBs in Arabidopsis (Wang *et al*, 2015). To test whether PPKs co-localized with RCD1 in the same NBs, we co-expressed RCD1-Venus and PPK-RFP transiently in tobacco. Results shown in **Figure 4A** demonstrated co-localization of RCD1 with all four RFP-tagged PPKs in NBs, but not with RFP alone. Expression of PPK-RFP constructs alone in tobacco showed uniform distribution of the proteins inside the nucleus (**Figure 4B**), which suggests that localization of Arabidopsis PPKs to NBs in tobacco was dependent on the interaction with Arabidopsis RCD1-Venus. Transient expression of PPK1-RFP, PPK3-RFP and PPK4-RFP in RCD1-Venus seedlings confirmed co-localization of RCD1 and PPKs in Arabidopsis (**Figure 4C**).

**Figure 4.**
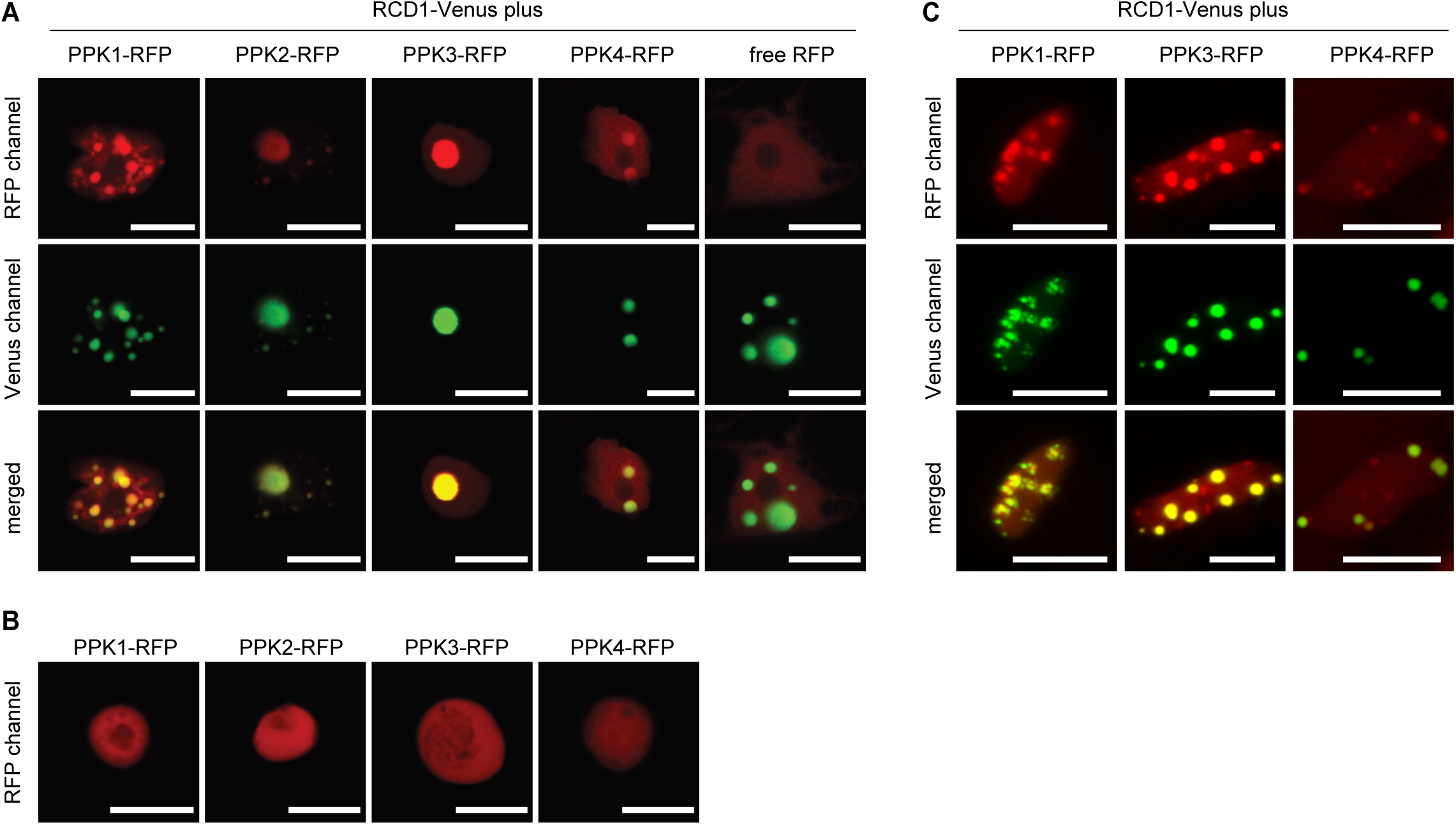
PPKs and RCD1 co-localize in NB. **A.** RCD1 co-localizes with PPKs in NBs in tobacco. RCD1-Venus was co-expressed with RFP or RFP-tagged PPKs in epidermal cells of *N. benthamiana* and the subnuclear localization was analyzed by confocal microscopy. Scale bars indicate 10 µm. **B.** PPK-RFPs alone do not form NBs when transiently expressed in tobacco. PPK-RFP fusion proteins were expressed as in (A) but without co-expression of RCD1-Venus. **C.** RCD1 co-localizes with PPK1, PPK3 and PPK4 in NBs in Arabidopsis. RFP-tagged PPKs were transiently expressed in Arabidopsis seedlings expressing RCD1-Venus and the subnuclear localization was analyzed by confocal microscopy. Scale bars indicate 10 µm.

These results are in line with complex formation between RCD1 and PPKs and confirmed their co-localization in NBs.

### RCD1 is phosphorylated by PPKs

Interaction of RCD1 with PPKs prompted us to study phosphorylation of RCD1 in more detail. Mass spectrometric determination of *in vivo* phosphosites in RCD1-HA immunoprecipitated from Arabidopsis revealed several phospho-serine and phospho-threonine-containing peptides (**Table 1, Figure 5A**). To verify whether PPKs could phosphorylate RCD1 directly, we tested PPK kinase activity towards RCD1 *in vitro* using recombinant GST-tagged proteins. GST-PPK2 and GST-PPK4 could be purified from *E. coli* with detectable kinase activity against the generic substrates myelin basic protein (MBP) and casein (**Supplementary figure 6**). Phosphorylation experiments using radioactively labelled γ[^32^P]-ATP (**Figure 5B, C**) showed that both GST-PPKs phosphorylated GST-RCD1 *in vitro*. Phosphorylated GST-RCD1 was analyzed by mass spectrometry to identify PPK-dependent *in vitro* phosphorylation sites. This revealed that several of the PPK-dependent *in vitro* phosphopeptides of RCD1 were also identified in the *in vivo* pull-down experiments. All *in vivo* and *in vitro* phosphopeptides from this and an earlier study (Wirthmueller *et al*, 2018) are listed in **Table 1**. A schematic representation of all identified phospho-sites in RCD1 is shown in **Figure 5A**. Notably, RCD1 is phosphorylated almost exclusively in its IDRs.

**Table 1:**
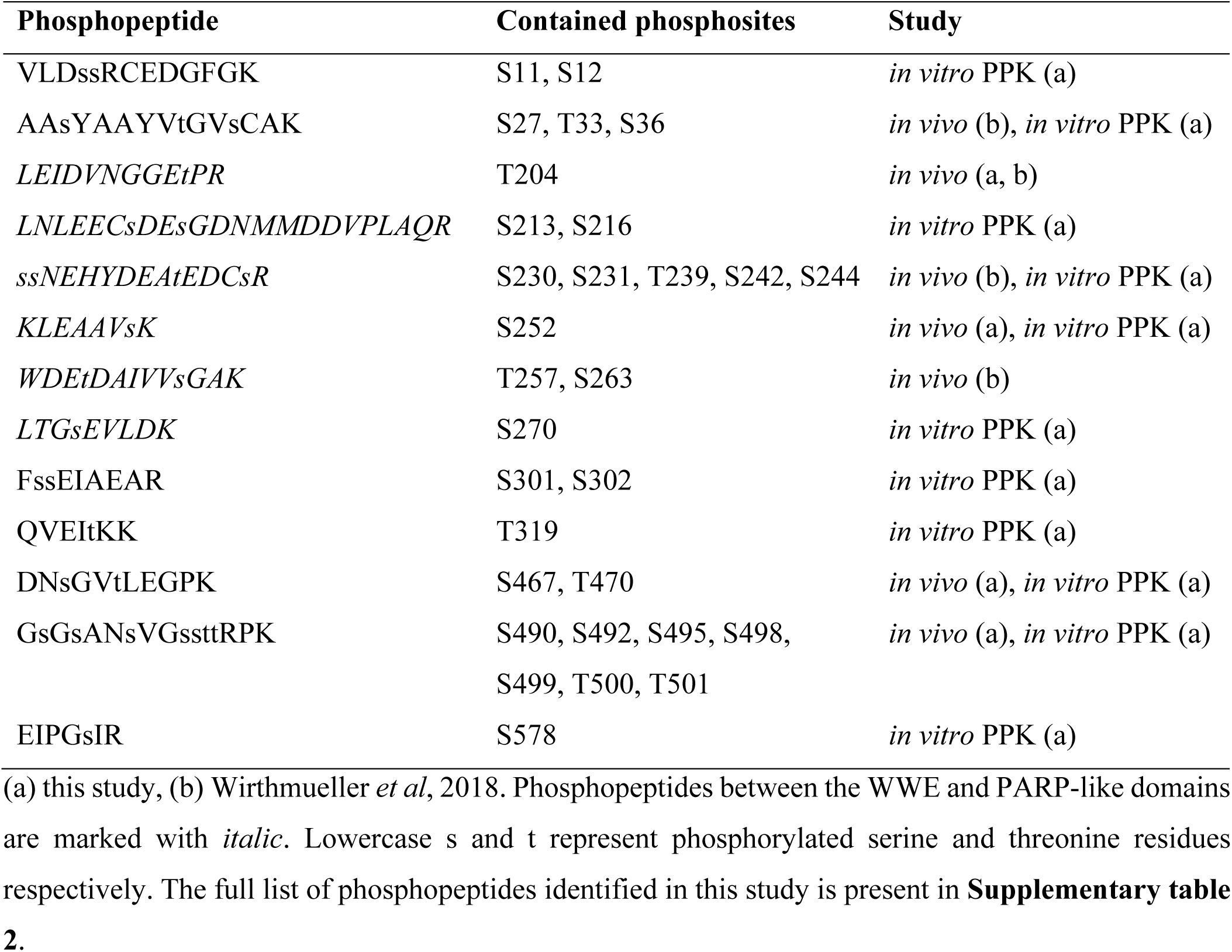
List of phosphosites identified *in vivo* or after *in vitro* kinase assay using PPKs.

**Figure 5.**
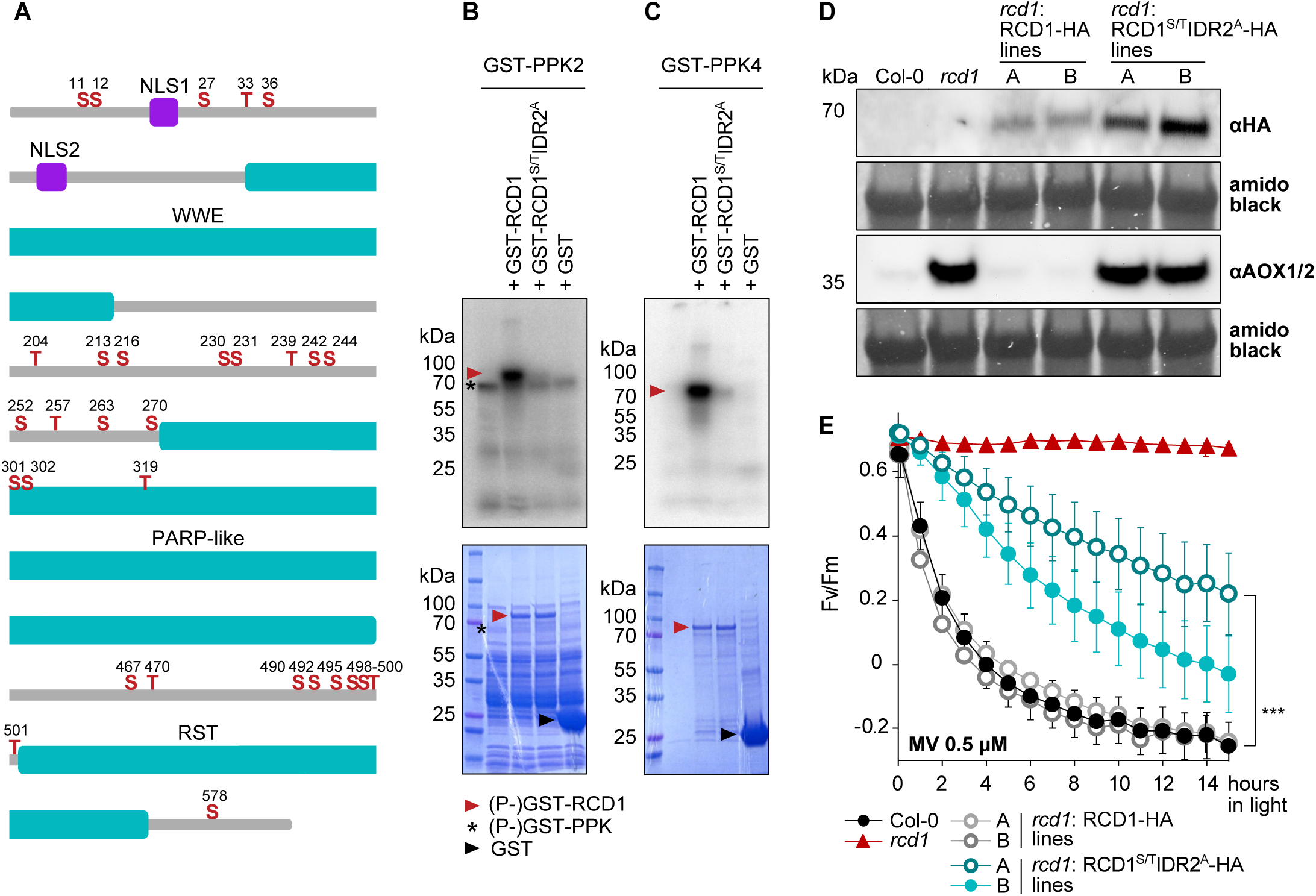
PPKs phosphorylate RCD1 at multiple sites. **A.** RCD1 phosphosites identified by *in vivo* and *in vitro* analyses as described in **Table 1**. RCD1 domains are highlighted in blue. Individual phosphosites are marked in red and numbered. **B, C.** Phosphosites in the region between WWE and PARP-like domains are targets for PPKs. Recombinant GST-PPK2 and GST-PPK4 were used together with recombinant GST-RCD1 protein in *in vitro* kinase assays. GST-PPK2 and 4 (asterisks) showed activity towards GST-RCD1 (red arrowhead). There was much less activity detected against the mutated GST-RCD1^S/T^IDR2^A^ protein. Upper panel shows autoradiograph, lower panel shows the Coomassie-stained SDS-PAGE. **D.** Phosphorylation of RCD1 IDR2 by PPKs affects its stability and function. *In vivo* abundance of RCD1^S/T^IDR2^A^-HA and of wild-type RCD1-HA variants was assessed in independent transgenic lines by immunoblot analysis with HA-specific antibody. The RCD1^S/T^IDR2^A^-HA variant did not fully complement *rcd1*-specific accumulation of alternative oxidases, as revealed by immunoblot with αAOX1/2 antibodies. Rubisco large subunit detected by amido black staining is shown as a control for equal protein loading. **E.** RCD1^S/T^IDR2^A^-HA variant does not fully complement *rcd1*-specific tolerance to MV. PSII inhibition (Fv/Fm) by MV was measured in indicated lines using 0.5 μM MV. For each experiment, leaf discs from at least four individual rosettes were used. The experiment was performed three times with similar results. Mean ± SD are shown. *** – P-value < 0.001 with Tukey corrected *post hoc* test at the selected time point between *rcd1*: RCD1^S/T^IDR2^A^-HA (line A) and *rcd1*: RCD1-HA (line A) lines. Full source data and statistics are presented in **Supplementary table 1**.

Combined data of *in vivo* and *in vitro* analyses of RCD1 phosphorylation revealed that most phosphosites concentrated in the IDR2, the region between the WWE and PARP-like domains. We mutated the 15 identified phosphosites in this region to non-phosphorylatable alanine residues by gene synthesis yielding an RCD1 variant that is further referred to as RCD1^S/T^IDR2^A^. This protein variant was subjected to *in vitro* kinase assays with GST-PPK2 and GST-PPK4. Mutation of the 15 phosphosites in IDR2 abolished phosphorylation of RCD1 by PPKs *in vitro* **(Figure 5B, C)**. Thus, PPKs showed specificity towards RCD1 phosphosites in IDR2. This is consistent with the previous result that the sequence up to the PARP-like domain was sufficient to co-immunoprecipitate endogenous PPKs from plant cell extracts (Wirthmueller *et al*, 2018). To address the role of IDR2 phosphorylation *in vivo,* we generated transgenic lines expressing the RCD1^S/T^IDR2^A^-HA construct in *rcd1* background under the native RCD1 promoter. In accordance with the *in-vitro* data, mutation of the IDR2 phosphosites to alanine resulted in disappearance of the phosphorylated protein form in *rcd1*: RCD1^S/T^IDR2^A^-HA line (**Supplementary figure 7A**). Despite the higher abundance, the RCD1^S/T^IDR2^A^-HA variant did not complement accumulation of AOX1/2 specific for *rcd1*, as revealed by immunoblot with αAOX1/2 antibodies (**Figure 5D**). Furthermore, expression of RCD1^S/T^IDR2^A^-HA could not fully complement the *rcd1* MV tolerance (**Figure 5E**), nor did it complement the habitus (**Supplementary figure 7B**) and the early flowering of *rcd1* (**Supplementary figure 2B**). This suggests that mutation of the 15 residues in IDR2 affected the nuclear functions of RCD1. Since the RST domain, which interacts with transcription factors, is retained in this mutant variant, phosphorylation by PPKs seems to be required for RCD1 to act as a negative regulator of transcription factors as proposed earlier (Jaspers et al, 2009; Vainonen et al, 2012; Shapiguzov et al, 2019).

To test the effect of RCD1 phosphorylation by PPKs on PAR binding ability *in vitro*, we performed dot-blot assay using purified non-phosphorylatable GST-RCD1^S/T^IDR2^A^ and phospho-mimicking GST-RCD1^S/T^IDR2^D/E^. Neither of the variants showed significantly different efficiency in binding purified PAR *in-vitro* as compared to wild type GST-RCD1 (**Supplementary figure 7C**). Thus, phosphorylation of IDR2 did not affect the interaction between RCD1 and purified PAR.

In addition to the PPK-related phosphosites between the WWE and the PARP-like domains, RCD1 harbors other *in vivo* phosphorylation sites, presumably targeted by other protein kinases linking RCD1 to different upstream signaling pathways **(Figure 5A)**. One of the identified sites, Thr204, is a predicted target for proline-directed protein kinases. We tested Arabidopsis GSK3/Shaggy-like protein kinases (ASKs), a group of stress-related proline-directed protein kinases (Saidi *et al*, 2012), for their ability to phosphorylate RCD1. *In vitro* kinase assays with several ASKs (**Supplementary figure 8A**) showed that ASKα, ASKɣ, and to a lesser extent, ASKε phosphorylated RCD1. Since Thr204 was the only phosphorylated residue flanked by a proline, we mutated Thr204 to alanine (RCD1-T204A). This mutation abolished or strongly reduced phosphorylation of RCD1 by the ASKs (**Supplementary figure 8B**) indicating that ASKα, ASKɣ, and ASKε target Thr204. The kinases targeting phospho-sites towards the N- and C-termini of RCD1 remain to be identified.

Our results with *rcd1*:RCD1^S/T^IDR2^A^-HA lines (**Figure 5D** and **Supplementary figure 7**) suggest that phosphorylation by PPKs affects the stability of RCD1. PPK-mediated phosphorylation of proteins has been shown to impact protein stability by targeting proteins for degradation (Ni *et al*, 2016; Liu *et al*, 2017; Chen *et al*, 2018). To address this question further, we analyzed RCD1 levels in triple *ppk* mutant plants. Immunoblot analysis of *ppk124* and *ppk234* with an RCD1-specific antibody revealed increased RCD1 levels compared to wild type plants (**Figure 6A**). Furthermore, in accordance with earlier results (Shapiguzov *et al*, 2019), higher accumulation of native RCD1, such as in triple *ppk* mutants, coincided with lower resistance of plants to MV compared to wild type (**Figure 6B**). These data suggest that PPK-dependent phosphorylation of RCD1 plays an important regulatory role for RCD1 protein stability and function.

**Figure 6.**
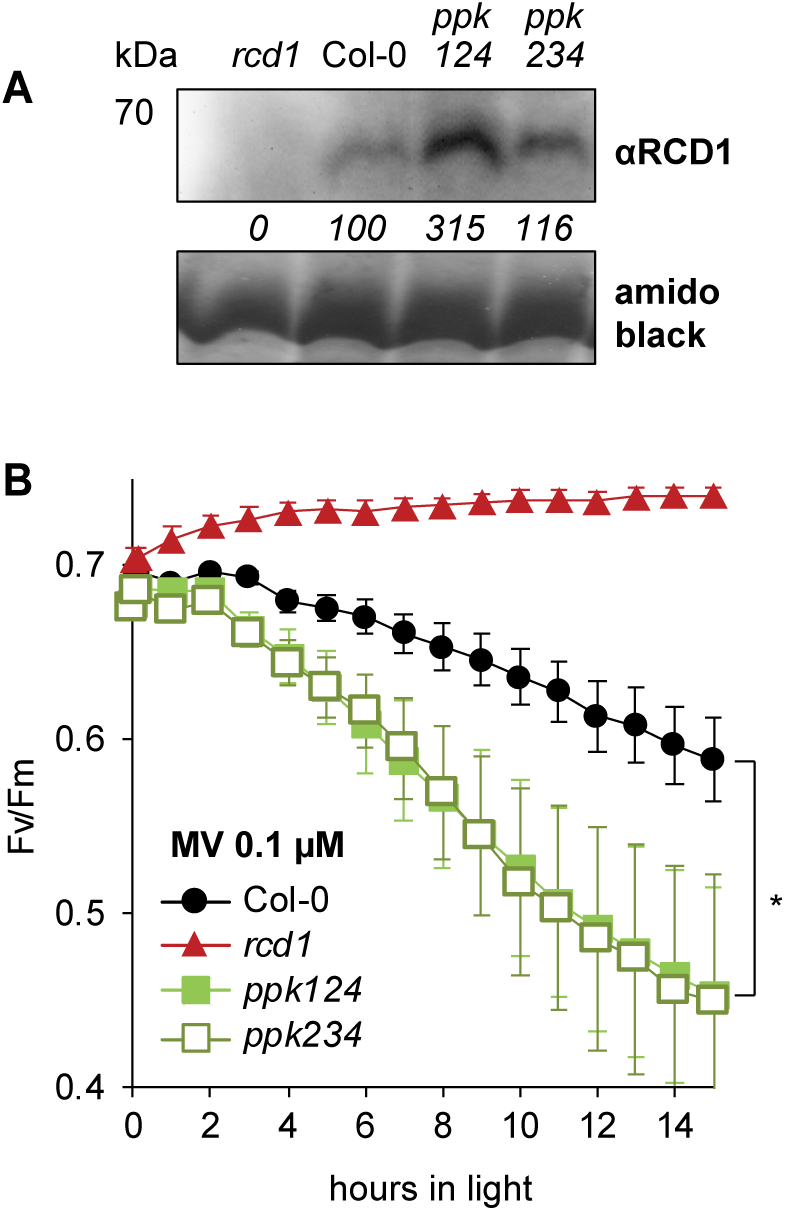
Knockout of PPKs stabilizes native RCD1. **A.** RCD1 accumulation in *ppk* triple mutants is higher than in Col-0. RCD1 level was assessed in total protein extracts from 3-week-old plants by immunoblot analysis with RCD1-specific antibody. The signal was quantified using ImageJ. The abundance in percent relative to Col-0 (100%) is shown under the immunoblot panel. Rubisco large subunit detected by amido black staining is shown as a control for equal protein loading. **B.** *ppk* triple mutants are more sensitive to MV than Col-0. PSII inhibition (Fv/Fm) by MV was measured in indicated lines using 0.1 μM MV. For each experiment, leaf discs from four individual rosettes were used. The experiment was performed three times with similar results. Mean ± SD are shown. * – P-value < 0.05 with Tukey corrected *post hoc* test at the selected time point between *ppk124* and Col-0. Source data and statistics are presented in **Supplementary table 1**.

## DISCUSSION

Plants continuously reprogram their gene expression in response to environmental stimuli. In nature, numerous simultaneous signals and cues have to be processed and integrated to achieve an adequate and balanced response. This can be accomplished e.g. with hub proteins which integrate signals from different sources and adjust the activity of transcription factors to ensure appropriate responses (Bugge *et al*, 2018; Vandereyken *et al*, 2018; Jespersen & Barbar, 2020). Hub proteins frequently interact with many different protein partners, including transcription factors, to provide a flexible system, which can simultaneously adjust several cellular functions according to changes in the surrounding environment. The RCD1 protein has been suggested in several studies to be such a hub protein (Jaspers *et al*, 2009; Hiltscher *et al*, 2014, Bugge *et al*, 2018; Shapiguzov *et al*, 2019; Jespersen & Barbar, 2020). Accordingly, disruption of the *RCD1* gene results in highly pleiotropic phenotypes and altered expression of a large number of genes (Ahlfors *et al*, 2004; Jaspers *et al*, 2009; Teotia & Lamb 2009, Brosché *et al*, 2014). Interaction of RCD1 with such a great variety of proteins is facilitated by its IDRs, which enable RCD1 to adjust its final conformation upon binding to its interaction partners (Kragelund *et al*, 2012; Bugge *et al*, 2018). In addition, other factors, such as recognition of signaling molecules or regulation of protein stability can contribute to the versatility of hub proteins, including RCD1.

### RCD1 is a PAR reader

Protein PARylation is a transient post-translational modification that has been associated with adjustment of development and response to stress conditions in plants (Briggs & Bent, 2011; Lamb *et al*, 2012; Feng *et al*, 2015). While the inventories of PARPs and PARGs have been defined in Arabidopsis (Vainonen *et al*, 2016; Rissel & Peiter, 2019), only a very limited number of PARylated proteins have so far been identified in plants; PARPs (Babiychuk *et al*, 1998; Feng *et al*, 2015), histones (Whitby *et al*, 1979; Willmitzer 1979) and the nuclear protein DAWDLE involved in micro-RNA processing (Feng *et al*, 2016). Nuclear Cajal bodies have also been linked to active PARPs in plants (Love *et al*, 2017).

The so-called “PAR readers”, proteins that non-covalently bind PAR (Teloni & Altmeyer, 2016; Gupte *et al*, 2017) have thus far remained unidentified in plants. Several animal WWE domain containing proteins have been described as PAR readers. However, it is worth to note that merely the presence of WWE domain in a protein sequence *per se* does not guarantee PAR binding, for instance the WWE domains of human PARP14 and DDHD2 are unable to bind PAR (Wang *et al*, 2012; He *et al*, 2012). It has been suggested that in animal cells the PAR polymer provides an interaction platform for PAR reader proteins to modulate cellular responses, including chromatin remodeling, protein degradation and cell death (Kim *et al*, 2020). In mammalian cells, several of these PAR-related processes co-localize with PAR-binding proteins in NBs (Ahel *et al*, 2008). The localization of RCD1 in NBs reported in this study was suppressed by 3MB (**Figure 3C**), a nicotinamide analog that inhibits PARP activity, suggesting that localization of RCD1 to NBs was PAR-dependent. However, the localization of RCD1 to NBs was also compromised if either the WWE or the PARP-like domain was removed, while *in vitro* experiments showed that only the WWE domain of RCD1 directly bound PAR. This suggests that the role of the PARP-like domain of RCD1 in its PAR-dependent NB localization may be related to other processes or interactions. Thus, the detailed molecular mechanisms whereby PAR participates in the formation of NBs and the recruitment of RCD1 therein remain to be elucidated.

While the WWE-PARP module is required for the PAR-dependent localization of RCD1 to NBs, the C-terminal RST domain of RCD1 binds many different transcription factors (Jaspers *et al*, 2009, 2010). Based on our data presented here and in previous publications, RCD1 might act as a negative regulator interacting with transcription factors in specific location in the nucleus or the chromatin and preventing their action by, for example, targeting them for degradation. Recognition of PARylated proteins by RCD1 could serve a scaffolding function, bringing together the components of the complex in NBs, including PARylated proteins, transcription factors, and regulatory protein kinases like PPKs.

Analysis of WWE domain proteins in animals has shown that the WWE domains co-exist in proteins not only with PARP/PARP-like domains, but also with E3 ubiquitin-ligase domains (Aravind 2001; Wang *et al*, 2012), which are involved in proteasomal degradation processes. Intriguingly, a significant fraction of transcription factors interacting with RCD1 is known to be regulated by proteasomal degradation (Qin *et al*, 2008; Ni *et al*, 2017; Favero *et al*, 2020). Furthermore, several proteins related to ubiquitin-dependent protein catabolism co-immunoprecipitated with RCD1 (Shapiguzov *et al*, 2019) and gene ontology analysis of altered gene expression in the *rcd1* mutant revealed enrichment in ubiquitin-proteasome-pathway associated genes (Jaspers *et al*, 2009). This supports a functional link between RCD1 and the nuclear proteasomal apparatus in Arabidopsis and suggests an evolutionary conserved link of PARylation and PAR readers with proteasomal degradation.

### Phosphorylation affects RCD1 stability and function

Hub proteins are often targets for multiple regulatory modifications enabling their involvement in a variety of upstream signaling processes. For example, RCD1 has recently been shown to perceive signals from organelles through thiol redox relays (Shapiguzov *et al*, 2019). Protein phosphorylation is another example of a ubiquitous post-translational protein modification that plays a major role in almost all physiological processes (Mergner *et al*, 2020). RCD1 is an *in vivo* phosphoprotein; overall, 13 phospho-peptides harboring potentially 30 phosphosites have been identified after immunoprecipitation of RCD1 from protein extracts. Notably, these phosphosites were enriched in the IDRs at the N-terminus as well as between the WWE, PARP-like, and RST domains of RCD1 (**Figure 5A**) and phosphorylation of these IDRs may have different function and outcome.

It has been shown that protein kinases preferentially target IDRs, and that phosphorylation can trigger disorder-to-order transitions of the protein structure (Iakoucheva *et al*, 2004; Bah & Forman-Kay, 2016). Structural analysis of RCD1 *in vitro* has shown that the disordered parts of the RST domain adapt their final folding only upon interaction with different transcription factors (Bugge *et al*, 2018; Shapiguzov *et al*, 2019). Additional phosphorylation of IDRs flanking the RST domain (IDR3 and IDR4) may influence the structure of RCD1 and its interaction with transcription factors *in vivo*.

In addition to the C-terminal phosphopeptides flanking the RST domain, we identified a phosphorylation hotspot in IDR2 between the WWE and the PARP-like domains targeted by PPKs. The IDR2 has recently been shown to be important for homo- or heterodimerization of RCD1 and its closest homolog SRO1 (Wirthmüller *et al*, 2018). Consequently, phosphorylation of IDR2 may affect the overall scaffolding structure of RCD1 and therefore regulate a wide variety of protein-protein interactions. Lack of phosphorylation of IDR2 in the RCD1^S/T^IDR2^A^ variant prevented full complementation of the *rcd1* phenotypes; however, it did not affect PAR binding *in-vitro*. It might be that the protein tertiary structure changed so that it cannot bind transcription factors. More likely, however, the RCD1^S/T^IDR2^A^ variant might still interact with transcription factors via the unaffected C-terminal RST domain. In this scenario, phosphorylation by PPKs would be necessary for targeting RCD1 and its partner transcription factors for degradation, and thus for the function of RCD1 as a negative transcriptional co-regulator. Supporting this mode of action: the activity of PPK4 delays flowering (Kang *et al,* 2020), while RCD1^S/T^IDR2^A^ results in early flowering similarly to *rcd1*. Accordingly the overexpression of the RCD1 interacting transcription factors BBX24 and FBH3 (Jaspers *et al*, 2009; Jaspers *et al*, 2010) has been shown to induce early flowering (Li *et al*, 2014; Ito *et* al, 2012). Thus in wild-type plants, these transcription factors may complex with RCD1 and after IDR2 gets phosphorylated by PPK4, be targeted for proteasomal degradation and thereby preventing overaccumulation of BBX24 and FBH3 that would cause early flowering.

IDR2 has also been reported to be required for the interaction between RCD1 and the oomycete effector protein HaRxL106 that prevents activation of plant immunity (Wirthmüller *et al*, 2018). Similarly, a *Phytophthora* RxLR effector has been shown to prevent relocalization of two tobacco NAC transcription factors from the endoplasmic reticulum to nucleus, which promoted disease progression (McLellan *et al*, 2013). Interestingly these tobacco NAC transcription factors, NTP1 and NTP2, are homologs of Arabidopsis ANAC013 and ANAC017, which are negatively regulated by RCD1 (Shapiguzov *et al,* 2019). Furthermore, it has been shown that ASKα, a potential upstream kinase of RCD1, has been contributing to plant immunity by modulating the oxidative pentose phosphate pathway (Stampfl *et al*, 2016). These results link RCD1 to the regulation of plant immunity and the phosphorylation of IDR2 appears to be involved in these processes.

Our results suggest that phosphorylation of RCD1 by PPKs regulates RCD1 stability. This is supported by the higher abundance of endogenous RCD1 in *ppk124* and *ppk234* triple mutants (**Figure 6A**). Accordingly, the *ppk124* and *ppk234* triple mutants exhibited increased sensitivity to MV as compared to wild type. Interestingly expression of the RCD1^S/T^IDR2^A^ form in *rcd1* led to only partial complementation of *rcd1* MV tolerance. This suggests that phosphorylation of IDR2 may not only be essential for RCD1 turnover alone or in complex with transcription factors but also for its function as transcriptional co-regulator.

Phosphorylation of other proteins by PPKs has been shown to alter protein stability (Liu *et al*, 2017; Ni *et al*, 2017; Chen *et al*, 2018). Ni *et al* (2017) described that PPKs phosphorylate the phyB-PIF3 complex upon light exposure, thereby targeting it for degradation - with an additional unknown factor X involved in the process. Intriguingly, RCD1 interacts with several PIFs, including PIF3 (Jaspers *et al*, 2009), which also localize to NBs (Favero, 2020). Moreover, PIF3 has been connected to retrograde signaling from the chloroplast (Martin *et al*., 2016), a process where RCD1 also plays a role (Shapiguzov *et al*, 2019).

The data presented here suggest a mechanism by which RCD1 levels could be regulated *via* phosphorylation by PPKs. This represents posttranslational control of a negative transcriptional co-regulator. Such regulation would allow PPKs to adjust the functions of RCD1 in response to environmental stimuli.

### Conclusions

Taken together, here we have provided experimental evidence for RCD1 as a PAR reader protein specific to plants. Our biochemical analyses revealed that the Arabidopsis RCD1 binds PAR and that the interaction is mediated by the WWE domain. The experimental data also suggests that RCD1 functions as a PAR-binding protein *in vivo*, making it a novel PAR reader protein described in plants. Furthermore, our results unveil a complex regulation of RCD1 function on post-translational level (**Figure 7**). RCD1 is targeted by its bipartite N-terminal NLS sequence to the nucleus (#1 in Figure 7) where it interacts with various proteins, including PPKs (#2) and transcription factors (#3), and accumulates in a PAR-dependent manner in NBs of unknown nature and composition (#4). Localization of RCD1 to NBs is mediated by its WWE and PARP domains. The C-terminal RST domain interacts with transcription factors (#3 in Fig. 7; Jaspers *et al*, 2009, 2010; Bugge *et al*, 2018), but whether the transcription factor is attached to the RST domain when PPKs phosphorylate IDR2 (#5 in Figure 7) is not yet known. Similarly, the exact order of the events #2, #3, and #4 is undetermined and the several possibilities require further studies.

**Figure 7.**
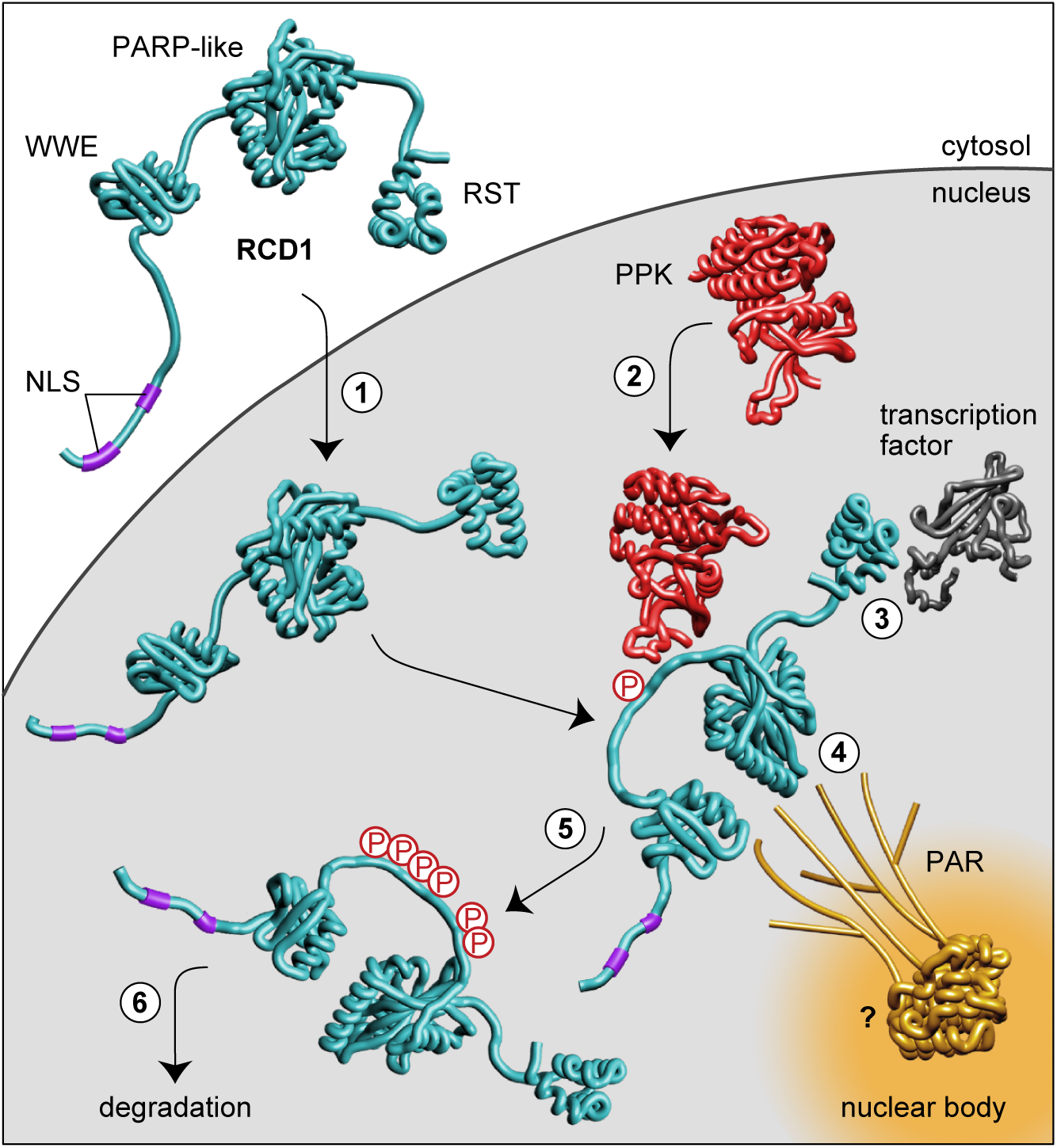
A model describing the regulation of nuclear RCD1 function in dependence of PAR binding and phosphorylation by PPKs. (1) RCD1 enters the nucleus by means of its bipartite N-terminal NLS sequence. In the nucleus, RCD1 interacts with PPKs (2), with diverse transcription factors (3) and with PAR (4). PAR recruits RCD1 to NBs of yet uncharacterized nature. Unknown PARylated proteins involved in RCD1 recruitment are labeled with a question mark. RCD1 is phosphorylated by PPKs at multiple sites in IDR2 (5), which targets RCD1 for degradation (6). RCD1 structure was predicted in RaptorX (http://raptorx.uchicago.edu/). Structural model of the WWE domain is based on mouse RNF146 (2RSF), structures of RCD1 PARP-like (5NGO, Wirthmueller *et al*, 2018) and RST (5N9Q, Bugge *et al*, 2018) domains have been reported. Terminal and inter-domain regions of RCD1 are not drawn to scale.

The ability of RCD1 to interact with a large number of transcription factors supports a function as a hub protein, which integrates various developmental as well as environmental signals. RCD1 might recognize PARylated proteins along the chromatin or in specific sub-nuclear domains and form complexes with transcription factors. Interaction with regulatory protein kinases PPKs and recruiting them to NBs would be a possible way of regulation of RCD1 function and stability of RCD1 and probably interacting transcription factors. Taken together, according to the data presented here, RCD1 represents the first described nuclear PAR-reader in plants. Therefore, our model proposes a new mechanism of fine-tuning transcriptional regulation, involving PAR-dependent compartmentalization and post-translational modification of the PAR-reader RCD1.

## MATERIALS AND METHODS

### Plants, mutants and chemical treatments

*Arabidopsis thaliana* plants were grown on soil (peat: vermiculite = 1:1) under white luminescent light (220-250 µmol m^-2^ s^-1^) at a 12-hour photoperiod and 22/18 °C. Seedlings were grown for 10 days on 1 x MS basal medium (Sigma) with 0.5 % Phytagel (Sigma). Arabidopsis *rcd1-4* mutant (GK-229D11) was used as a background for all complementation lines. The *ppk* triple mutants were kindly provided by Dr Dmitri Nusinow (Donald Danforth Plant Science Center, St. Louis) and have been described in Huang *et al*, (2016). Treatments with chemicals methyl viologen (MV, 0.1, 0.5, or 1 μM, as indicated in the figures) and 3-methoxybenzamide (3MB, 10 mM) were performed on leaf discs floating on Milli-Q water supplemented with 0.05% Tween 20 (Sigma), overnight at room temperature or at 4°C, accordingly. For 4′,6-diamidino-2-phenylindole (DAPI) staining, the seedlings were vacuum-infiltrated with 0.1 mM DAPI in Milli-Q water supplemented with 0.05% Tween 20.

### Plasmids

Full-length AtRCD1, the WWE-domain (amino acids 1-155), RCD1ΔWWE (consisting of PARP- and RST-domains, amino acids 241-589), RCD1ΔPARP (missing the residues 304-443) and the C-terminal part of RCD1 including the RST-domain (amino acids 468-589), were cloned into pGEX4T-1 for N-terminal GST fusion using primers listed in **Supplementary table 3**. Full-length AtRCD1 was also cloned into the pET8c vector for N-terminal His-fusion (Jaspers *et al*, 2010). For generating N-terminal GST-fusion constructs, PPK1-4 cDNAs were cloned into pGEX6P-1, and ASK cDNAs into pGEX4T-1. The kinase-dead ASK loss-of-function constructs contain a Lys-Arg mutation in the kinase activation loop.

For generating a GST fusion construct of RCD1 where the IDR2 is non-phosphorylateable (GST-RCD1^S/T^IDR2^A^), all phospho-serine and phospho-threonine residues within IDR2 were mutated to alanine residues by gene synthesis (Genescript Biotech, Netherlands).

To generate the RCD1-Venus construct, RCD1 cDNA was fused to the *UBIQUITIN10* promoter region and to the C-terminal triple Venus YFP tag in a MultiSite Gateway reaction as described in Siligato *et al*, (2016). The ΔWWE (missing the residues 90-151), ΔPARP (missing the residues 304-443) and ΔRST (missing the residues 462-589) deletions were introduced by PCR using primers listed in **Supplementary table 3** and end-joining using In-Fusion (Clontech).

Construction of transgenic lines expressing HA-tagged RCD1 (RCD1-3xHA) is described in Jaspers *et al*, (2009). RCD1*nls*-HA variant was made using the vector pDONR/Zeo that contained the RCD1 promoter followed by the wild-type genomic RCD1 sequence (Jaspers *et al*, 2009). PCR was performed with Q5 High-Fidelity DNA Polymerase (New England Biolabs) and the primers listed in **Supplementary table 3**. After sequential mutation of the two parts of the bipartite NLS, the construct was transferred to the Gateway pGWB13 binary vector and introduced into the plants as described in Jaspers *et al*, (2009). The ΔWWE (missing the residues 90-151), ΔPARP (missing the residues 304-443) and ΔRST (missing the residues 462-589) deletions were introduced by PCR using primers listed in **Supplementary table 3** and end-joining using In-Fusion (Takara).

For generating epitope-tagged PPK fusions, the coding sequences of the four *PPK* genes lacking their stop codons were cloned into NcoI/XhoI-digested pENTR4 using In-Fusion enzyme (Takara). The *PPK* coding sequences were then recombined by Gateway® Clonase II reactions into pH7WGR2 (Karimi *et al*, 2002) or pGWB414 (Nakagawa *et al*, 2007) to create RFP and 3xHA-tagged variants, respectively.

### Spectroscopic measurements of photosynthesis

Chlorophyll fluorescence was measured by MAXI Imaging PAM (Walz, Germany) essentially as described in Shapiguzov *et al*, (2019). PSII photoinhibition protocol consisted of repetitive 1-hour periods of blue actinic light (450 nm, 80 µmol m^-2^ s^-1^) each followed by a 20-min dark adaptation, then Fo and Fm measurement. PSII photochemical yield was calculated as Fv/Fm = (Fm-Fo)/Fm. The assays were performed in 96-well plates. In each assay leaf discs from at least 4 individual plants were analyzed. Each assay was reproduced at least three times.

### SDS-PAGE and immunoblotting

For immunoblotting of total plant extracts, the plant material was frozen immediately after treatments in liquid nitrogen and ground. Total proteins were extracted in SDS extraction buffer (50 mM Tris, pH 7.8, 2 % SDS, 1 x protease inhibitor cocktail; P9599, Sigma), 2 mg/ mL NaF) for 20 min at 37°C and centrifuged at 18 000 x g for 10 min. Supernatants were normalized for protein concentration and resolved by SDS-PAGE. After electrophoresis, proteins were electroblotted to PVDF membrane and probed with specific antibodies: αHA (Roche), αGFP (Milteny Biotech), αGST (Sigma), αPAR (Trevigen), αRCD1 (Shapiguzov *et al*, 2019), and αAOX1/2 (Agrisera AS04 054). The signal was visualized by ECL Prime chemiluminescence reagents (GE Healthcare). Quantification of the signal was done using ImageJ.

### Confocal microscopy

The subcellular localization of RCD1 in stable expression Arabidopsis line was analyzed by confocal microscopy with a Leica SP5 II HCS inverted microscope using a solid-state blue laser was used for visualizing YFP and chloroplast autofluorescence (detection with 521–587 and 636– 674 nm range, respectively). For co-localization studies of RCD1-Venus and PPK-RFP fusion constructs, the binary plasmids were transformed into *A. tumefaciens* strain GV3101 pMP90. Proteins were transiently expressed in *N. benthamiana* leaves as described below for co-immunoprecipitation assays. YFP was excited using a 488 nm laser with a detection window of 519-556 nm and RFP was excited using a 561 nm laser with detection at 599-657 nm.

### Protein expression and purification

Fusion proteins were expressed in *E.coli* BL21 (DE3) Codon Plus strain and purified using GSH- or Ni^2+^-Sepharose beads (GE Heathcare) according to manufacturer instructions as described before (Jaspers *et al*, 2009; Jaspers *et al*, 2010). The N-terminal GST-tagged WWE-domain of RNF146 (amino acids 100-175) was expressed and purified as described in Zhang *et al*, (2011).

### Poly(ADP-ribose) dot-blot assay

Purified His and GST fusion proteins or GST alone (500 ng) were blotted onto nitrocellulose membrane (BioRad). The nitrocellulose membrane was rinsed with TBS-T buffer (10 mM Tris-HCl at pH 7.4, 150 mM NaCl and 0.05 % Tween 20) three times. The membrane was incubated with 100 nM of purified PAR (Trevigen, 4336-100-01, 10 µM stock, polymer size 2-300 units) for 1 h at room temperature. After 5 washes with TBS-T and TBS-T containing 1 M NaCl, the membrane was blocked with 5 % milk followed by immunoblotting with mouse αPAR (Trevigen) or αGST (Sigma) antibody.

### Surface plasmon resonance

Recombinant RCD1-His and GST-RCD1ΔWWE proteins were coupled to a Biacore CM5 sensor chip *via* amino-groups. PAR (625 nM) was profiled at a flow rate of 30 mL/min for 300 s, followed by 600 s flow of wash buffer (10 mM HEPES, pH 7.4, 150 mM NaCl, 3 mM EDTA, 0.05% Surfactant P20). Mono ADP-ribose and cyclic ADP-ribose were profiled at 1 mM concentration. After analysis in BiaEvalution (Biacore), the normalized resonance units were plotted over time with the assumption of one-to-one binding.

### Transient protein expression in N. benthamiana and Arabidopsis

Binary vectors harbouring RCD1-GFP or PPK-3xHA fusions were transformed into *A. tumefaciens* strain GV3101 pMP90. For expression, Agrobacteria were scraped from selective YEB plates and resuspended in infiltration medium (10 mM MES pH 5.6, 10 mM MgCl_2_) and the OD_600_ was adjusted to 0.8. To suppress transgene silencing, Agrobacteria expressing the tomato bushy stunt virus 19K silencing suppressor were co-infiltrated. After adding acetosyringone to a final concentration of 100 μM and incubation for 2 h at room temperature, Agrobacteria were mixed in a ratio of 1:1:2 (19K) and infiltrated into *N. benthamiana* leaves.

For transient Arabidopsis expression the FAST co-cultivation technique was used (Li *et al*, 2009). In short binary vectors harbouring PPK-RFP fusions were transformed into A, tumefaciens strain GV3101 pMP90. From overnight liquid LB-culture Agrobacteria were washed and resuspended in co-cultivation medium to OD_600_ 2.5. Seedlings grown for 5 days in long days (16/8, light/dark) were soaked in Agrobacteria containing co-cultivation medium for 20 minutes.

### Co-immunoprecipitation

Infiltrated leaf tissue was harvested 72 h later and proteins were extracted by grinding leaf tissue in liquid nitrogen followed by resuspension in extraction buffer (50 mM Tris-HCl pH 7.5, 150 mM NaCl, 10% Glycerol, 1 mM EDTA, 5 mM DTT, 1x protease inhibitor cocktail [P9599, Sigma], 10 μM MG132) at a ratio of 2 mL / g FW. Protein extracts were centrifuged at 20000 x g / 4°C/20 min and a fraction of the supernatant was saved as ‘input’ sample. 15 μL of αGFP-nanobody:Halo:His6 magnetic beads (Chen *et al*, 2018) were added to 1.5 mL of protein extract followed by incubation on a rotating wheel at 4°C for 5 min. The beads were washed 3 times with 1 mL extraction buffer using a magnetic tube rack and then boiled in 80 μL SDS sample buffer to elute protein from the beads. For immunoblots, protein samples were separated by SDS-PAGE and electro-blotted onto PVDF membrane. Antibodies used were αGFP (210-PS-1GP, Amsbio) and αHA (11867423001, Sigma).

### Kinase activity assays

*In vitro* kinase assays using recombinant proteins were performed in a total volume of 20 µL of kinase buffer (20 mM HEPES, pH 7.5, 15 mM MgCl_2_, and 5 mM EGTA). The reaction was started with 2 μCi [γ-^32^P]ATP and incubated at room temperature for 30 min. The reaction was stopped by the addition of 5 µL of 4x SDS loading buffer. Proteins were resolved by SDS-PAGE and the gel was dried and exposed overnight to a phosphor imager screen. For the kinase activity test, GST-PPKs were tested against 5 µg myelin basic protein (MBP; Sigma Aldrich) and 5 µg Casein in 0.1 M Tris pH 8.8 (Sigma). For identification of *in vitro* phosphorylation sites by LC-MS/MS, 1.5 mM unlabeled ATP was used in the kinase buffer. The proteins were separated by SDS-PAGE, followed by Coomassie Brilliant Blue staining and were digested by trypsin (Promega).

### LC-MS/MS

Phosphopeptides were enriched from tryptic digests using TiO_2_ microcolumns (GL Sciences Inc., Japan) as described in Larsen *et al,* (2005). Enriched phosphopeptides were analyzed by a Q Exactive mass spectrometer (Thermo Fisher Scientific) connected to Easy NanoLC 1000 (Thermo Fisher Scientific). Peptides were first loaded on a trapping column and subsequently separated inline on a 15-cm C18 column (75 μm × 15 cm, ReproSil-Pur 5 μm 200 Å C18-AQ, Dr. Maisch HPLC). The mobile phase consisted of water with 0.1% (v/v) formic acid (solvent A) or acetonitrile/water (80:20 [v/v]) with 0.1% (v/v) formic acid (solvent B). A linear 60-min gradient from 6 to 42% (v/v) B was used to elute peptides. Mass spectrometry data were acquired automatically by using Xcalibur 3.1 software (Thermo Fisher Scientific). An information-dependent acquisition method consisted of an Orbitrap mass spectrometry survey scan of mass range 300 to 2000 m/z (mass-to-charge ratio) followed by higher-energy collisional dissociation (HCD) fragmentation for 10 most intense peptide ions. Raw data were searched for protein identification by Proteome Discoverer (version 2.2) connected to in-house Mascot (v. 2.6.1) server. Phosphorylation site locations were validated using phosphoRS algorithm. A SwissProt database (https://www.uniprot.org/) was used with a taxonomy filter Arabidopsis. Two missed cleavages were allowed. Peptide mass tolerance ± 10 ppm and fragment mass tolerance ± 0.02 D were used. Carbamidomethyl (C) was set as a fixed modification and Met oxidation, acetylation of protein N-terminus, and phosphorylation of Ser and Thr were included as variable modifications. Only peptides with a false discovery rate of 0.01 were used.

## ACKNOWLEDGEMENTS

We thank Mr Damien Marchese and Dr. Melanie Carmody for the help in generating the transgenic lines. We thank Dr. Dmitri Nusinow for providing the seeds of *ppk* triple mutants. We thank Maria Aatonen and Maria Semenova for help in Biacore experiments which were performed at the Biomolecular Interaction Unit, Faculty of Biological and Environmental Sciences, University of Helsinki. We thank Mika Molin and Marko Crivaro for help with confocal microscopy at the Light Microscopy Unit of the Institute of Biotechnology, University of Helsinki. We thank Dr. Romy Schmidt for her input in studying the RCD1 nuclear bodies. Mass spectrometry analyses were performed at the Turku Proteomics Facility, supported by Biocenter Finland. This work was supported by the University of Helsinki (JK); the Academy of Finland Centre of Excellence programs (2006-11; and 2014-19; JK) and Research Grant (Decision 250336; JK). MW acknowledges funding from the Academy of Finland (Decisions 275632, 283139, 312498, and 323917). LW acknowledges funding by the German Research Foundation (DFG; grant WI 3670/2-1) and core funding from the Leibniz Institute of Plant Biochemistry (IPB) and the FU Berlin Dahlem Centre of Plant Sciences. RG acknowledges funding from the Doctoral Programme in Plant Sciences by the University of Helsinki.

## AUTHOR CONTRIBUTION

JV, AS, JKW, RG, CJ, LW, MW, and JK conceived and designed experiments. JV, AS, JKW, RG, RDM, ID, TP, NB, and LW carried out experiments. JV, AS, JKW, RG, RDM, ID, NB, CJ, LW, MW, and JK analyzed the data. JV, AS, JKW, RG, LW, and JK wrote the article. All authors read and contributed to the final article.

## Supplementary information

**Supplementary figure 1. RCD1 localization and the domain deletion constructs used in the study.**

**A.** RCD1*nls*-Venus is localized outside the nuclei. Confocal images were taken from stable Arabidopsis lines expressing full-length RCD1-Venus and RCD1*nls*-Venus. DAPI staining was used to highlight nuclear structures. White bars indicate 10 µm.

**B.** Schematic representation of RCD1 domain deletion constructs fused to triple HA or triple Venus tag and expressed in *rcd1* background.

**Supplementary figure 2. Characterization of stable transgenic lines expressing RCD1 domain deletion constructs fused to triple HA tag in *rcd1* background.**

**A.** MV sensitivity is not restored in lines expressing RCD1ΔWWE-HA (ΔW) and RCD1ΔPARP-HA (ΔP), and RCD1ΔRST-HA (ΔR). Two independent lines for each construct (A and B) were used in the experiments. PSII inhibition (Fv/Fm) by MV was measured in indicated lines using 0.5 μM MV. For each experiment, leaf discs from three individual rosettes were used. The experiment was performed three times with similar results. Source data and statistics are presented in **Supplementary** table 1.

**B.** Early flowering time phenotype of *rcd1* is not restored in lines expressing RCD1ΔWWE-HA (ΔW) and RCD1ΔPARP-like-HA (ΔP), and RCD1ΔRST-HA (ΔR). The RCD1^S/T^IDR2^A^-HA variant is addressed below. Two independent lines for each construct (A and B) were used in the experiments. Flowering time defined as the day of the opening of the first flower after germination, is plotted against the number of expanded rosette leaves on the flowering day. The experiment was performed three times with similar results. Source data and statistics are presented in **Supplementary** table 1.

**Supplementary figure 3. RCD1 binds PAR but not ADP-ribose or cyclic ADP-ribose.**

**A.** The purity of recombinant proteins used in *in vitro* analyses of PAR binding. Proteins were purified, resolved by SDS-PAGE and stained with Coomassie.

**B.** GST-RCD1ΔPARP-like binds PAR *in vitro*. PAR binding activity of immobilized GST-RCD1ΔPARP-like and GST-RCD1 was assessed by dot-blot assay using PAR-specific antibody. GST was used as a negative control and GST antibody was used to assess protein loading.

**C.** PAR titration curve obtained by SPR analysis of PAR binding by RCD1-His. The curve was plotted using non-linear regression with the assumption of one-to-one binding.

**D, E**. RCD1-His binds neither mono-ADP-ribose (ADPR), nor cyclic ADP-ribose (cADPR). SPR sensorgrams do not show any response in case of ADPR or cADPR profiled at 1 mM concentrations over immobilized RCD1-His.

**Supplementary figure 4. RCD1 is phosphorylated *in vivo*.** RCD1-HA migrates in SDS-PAGE as a double band, as visualized by immunoblot analysis of protein extracts with HA-specific antibody. Upper band corresponding to phosphorylated form of RCD1-HA was diminished by treatment of plant extracts with alkaline phosphatase (CIP). Rubisco large subunit detected by amido black staining is shown as a control for equal protein loading.

**Supplementary figure 5. RCD1-GFP interacts with PPK-HA in tobacco.** RCD1-GFP was transiently co-expressed with HA-tagged versions of PPK1, 2, 3 or 4 in *N. benthamiana*. YFP served as negative control. At 72 hours post infiltration, RCD1-GFP and YFP were immunoprecipitated with GFP-specific antibody and co-precipitating PPK-HA proteins were detected by αHA immunoblot. Immunoprecipitation of RCD1-GFP and YFP was confirmed by an αGFP immunoblot. ‘Input’ samples were taken before immunoprecipitation and included on the immunoblots to test for equal expression and loading.

**Supplementary figure 6. Recombinant PPK2 and PPK4 are active in *in vitro* kinase assays.** Recombinant GST-PPK1-4 were used together with generic substrates casein and myelin basic protein (MBP) in an *in vitro* kinase assay. Upper panel shows autoradiograph, lower panel shows the Coomassie-stained SDS-PAGE.

**Supplementary figure 7. Effect of RCD1 phosphorylation in IDR2 by PPKs on the protein function. A.** *In vivo* phosphorylation pattern of RCD1^S/T^IDR2^A^-HA is different from that of the wild type RCD1-HA. Upper band corresponding to phosphorylated form of RCD1 is not detectable in RCD1^S/T^IDR2^A^-HA line as visualized by immunoblot analysis of protein extracts with HA-specific antibody. The lines with approximately equal expression of RCD1 were selected for this comparison. 100% corresponds to 100 μg of total protein. Rubisco large subunit detected by amido black staining is shown as a control for equal protein loading.

**B.** Early flowering phenotype of *rcd1* is not restored by expression of RCD1^S/T^IDR2^A^-HA in *rcd1* background. Picture shows 5-week-old plants of representative lines under standard growth conditions. Domain deletion lines are shown in the same figure for comparison.

**C.** PPK phosphorylation in IDR2 does not affect PAR binding by RCD1*in vitro*. PAR binding activity of immobilized GST-tagged RCD1^S/T^IDR2^A^ and RCD1^S/T^IDR2^D/E^, mimicking non-phosphorylated or fully phosphorylated RCD1, respectively, was accessed by dot-blot assay using PAR-specific antibody. GST was used as a negative control. GST antibody was used to assess protein loading.

**Supplementary figure 8. ASKα, ASKγ, and ASKε phosphorylate RCD1 *in vitro*.**

**A.** Specificity of ASKα, ASKγ and ASKε towards RCD1. Recombinant ASK-GSTs were used together with recombinant GST-RCD1 protein in an *in vitro* kinase assay. P – phosphorylated protein; autoP – autophosphorylated protein.

**B.** Thr204 is the target for ASKs. ASKα,γ,ε-GST were used with recombinant GST-RCD1 or GST-RCD1T204A in an *in vitro* kinase assay. LOF indicates loss-of-function constructs of ASKs.

Upper panels show autoradiographs, lower panels show the Coomassie-stained SDS-PAGE.

**Supplementary table 1.** Source data and statistical analyses.

**Supplementary table 2.** Identified RCD1 phosphopeptides.

**Supplementary table 3.** Primers used in the study.

