## Supplemental figures 1-8 for "Arabidopsis poly(ADP-ribose)-binding protein RCD1 interacts with Photoregulatory Protein Kinases in nuclear bodies"

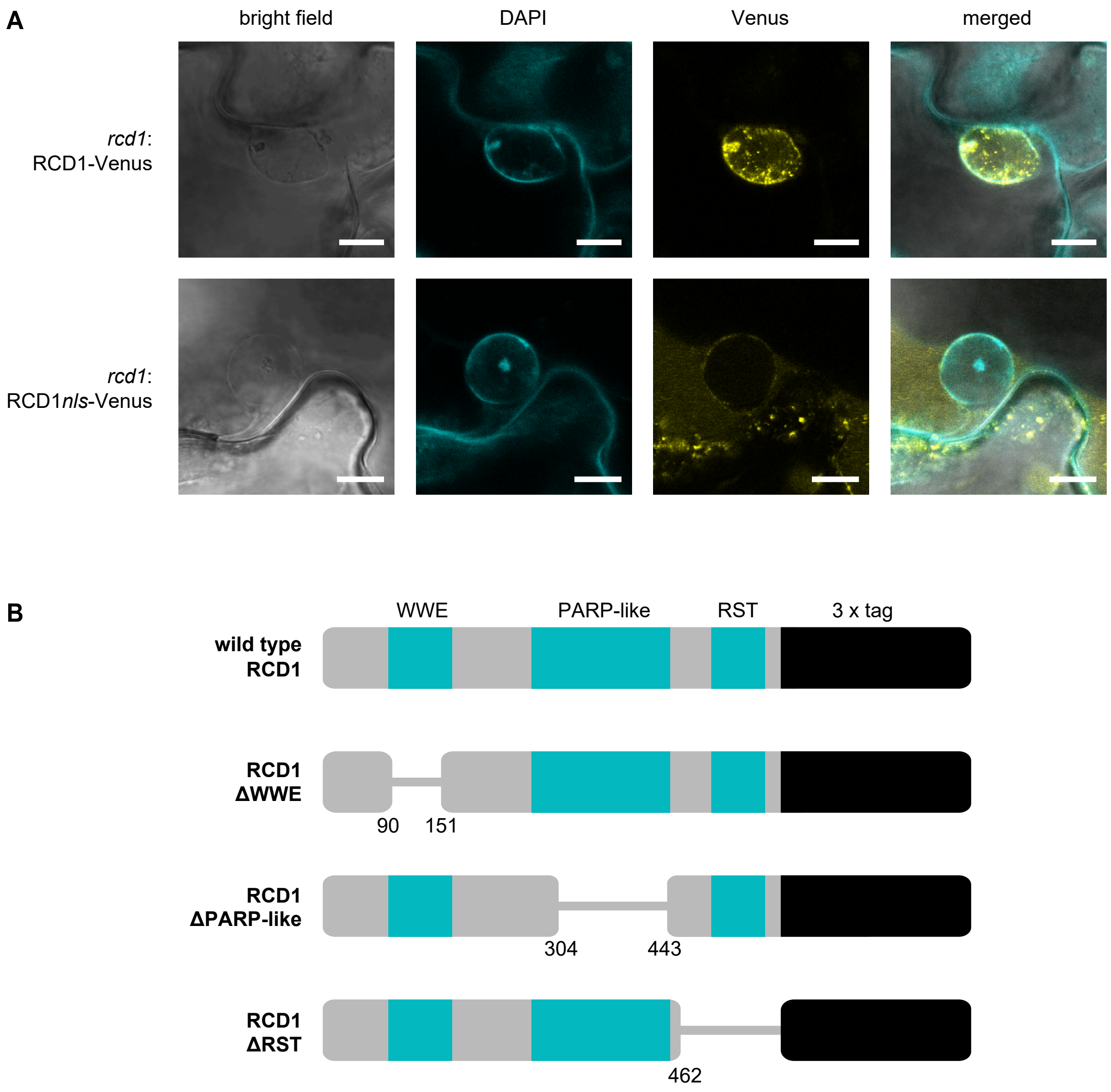


**Supplementary figure 1. RCD1 localization and the domain deletion constructs used in the study.**
**A.** RCD1*nls*-Venus is localized outside the nuclei. Confocal images were taken from stable Arabidopsis lines expressing full-length RCD1-Venus and RCD1*nls*-Venus. DAPI staining was used to highlight nuclear structures. White bars indicate 10 μm.
**B.** Schematic representation of RCD1 domain deletion constructs fused to triple HA or triple Venus tag and expressed in *rcd1* background.

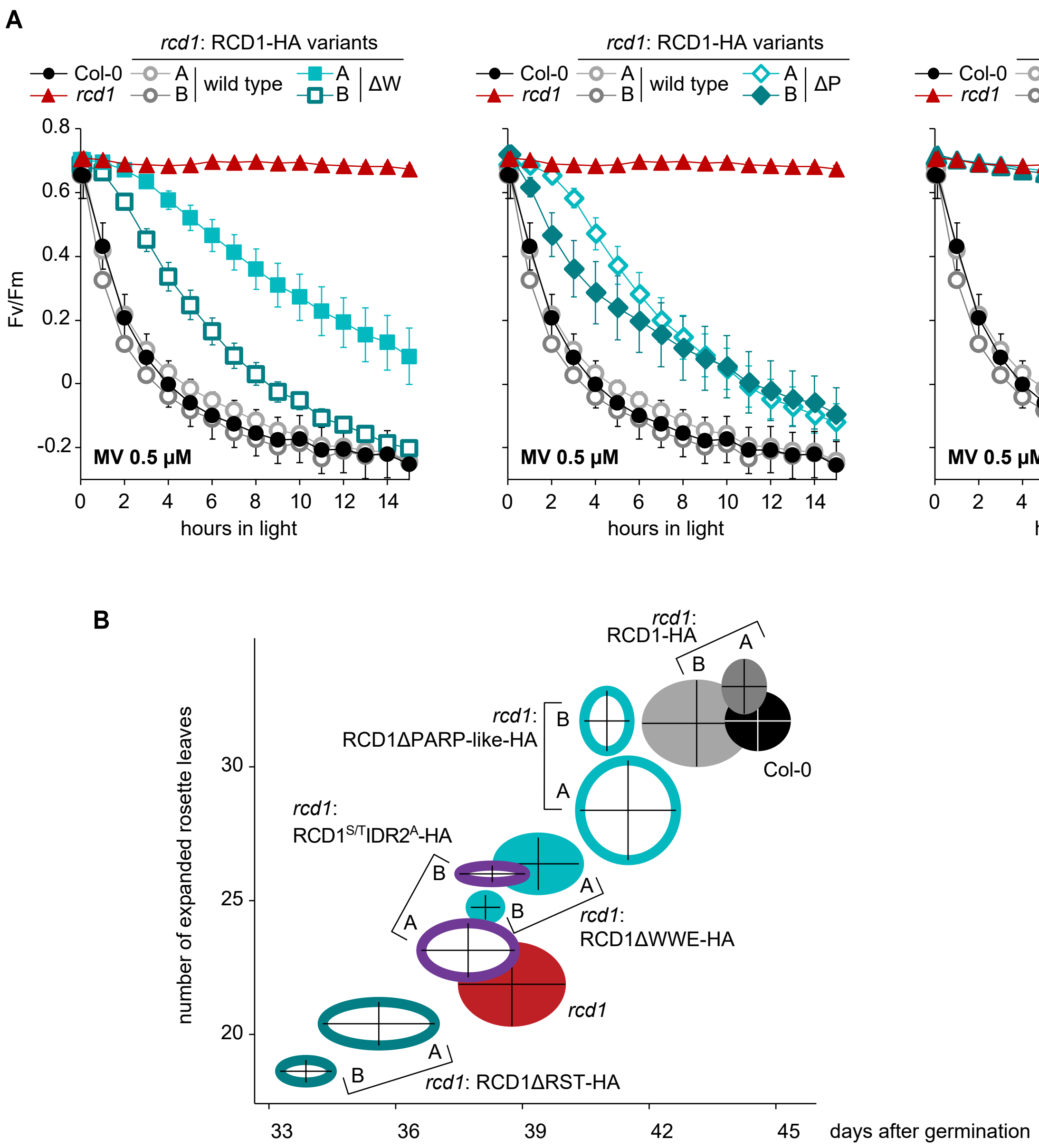


**Supplementary figure 2. Characterization of stable transgenic lines expressing RCD1 domain deletion constructs fused to triple HA tag in *rcd1* background.**

**A.** MV sensitivity is not restored in lines expressing RCD1ΔWWE-HA (ΔW) and RCD1ΔPARP-HA (ΔP), and RCD1ΔRST-HA (ΔR). Two independent lines for each construct (A and B) were used in the experiments. PSII inhibition (Fv/Fm) by MV was measured in indicated lines using 0.5 µM MV. For each experiment, leaf discs from three individual rosettes were used. The experiment was performed three times with similar results. Source data and statistics are presented in **Supplementary table 1**.

**B.** Early flowering time phenotype of *rcd1* is not restored in lines expressing RCD1ΔWWE-HA (ΔW), RCD1ΔPARP-like-HA (ΔP), and RCD1ΔRST-HA (ΔR). The RCD1$^{S/T}$IDR2$^{A}$-HA variant is addressed below. Two independent lines for each construct (A and B) were used in the experiments. Flowering time defined as the day of the opening of the first flower after germination, is plotted against the number of expanded rosette leaves on the flowering day. The experiment was performed three times with similar results. Source data and statistics are presented in **Supplementary table 1**.

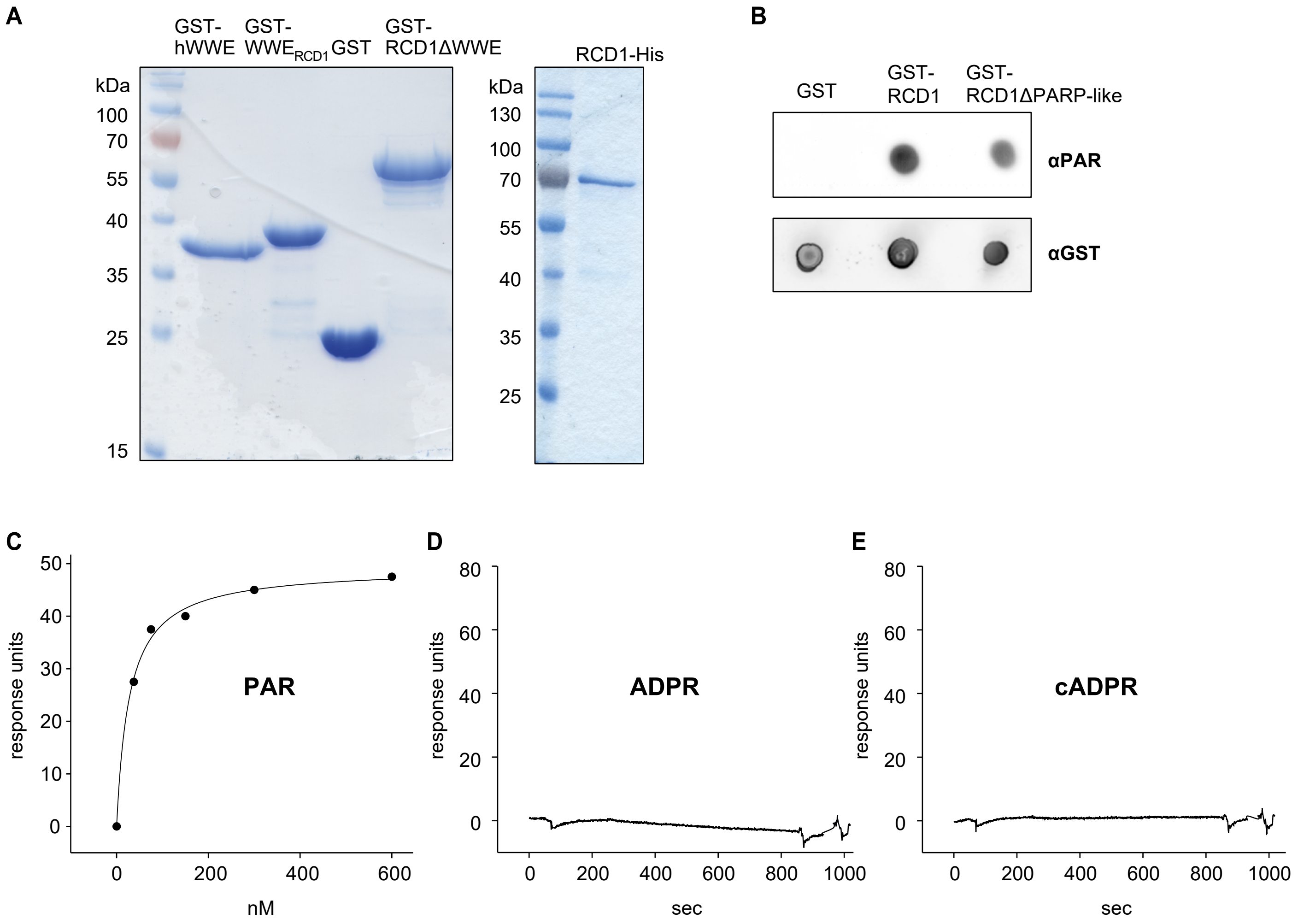


**Supplementary figure 3. RCD1 binds PAR but not ADP-ribose or cyclic ADP-ribose.**

**A.** The purity of recombinant proteins used in *in vitro* analyses of PAR binding. Proteins were purified, resolved by SDS-PAGE and stained with Coomassie.

**B.** GST-RCD1ΔPARP-like binds PAR *in vitro*. PAR binding activity of immobilized GST- RCD1ΔPARP-like and GST-RCD1 was assessed by dot-blot assay using PAR-specific antibody. GST was used as a negative control and GST antibody was used to assess protein loading.

**C.** PAR titration curve obtained by SPR analysis of PAR binding by RCD1-His. The curve was plotted using non-linear regression with the assumption of one-to-one binding.

**D, E.** RCD1-His binds neither mono-ADP-ribose (ADPR), nor cyclic ADP-ribose (cADPR). SPR sensorgrams do not show any response in case of ADPR or cADPR profiled at 1 mM concentrations over immobilized RCD1-His.

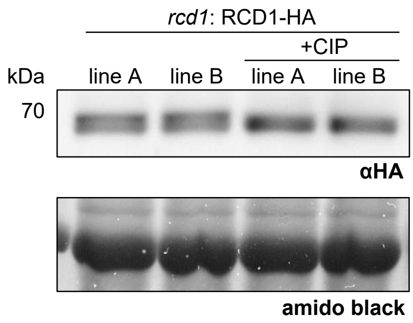


**Supplementary figure 4. RCD1 is phosphorylated *in vivo*.**

RCD1-HA migrates in SDS-PAGE as a double band, as visualized by immunoblot analysis of protein extracts with HA-specific antibody. Upper band corresponding to phosphorylated form of RCD1-HA was diminished by treatment of plant extracts with alkaline phosphatase (CIP). Rubisco large subunit detected by amido black staining is shown as a control for equal protein loading.

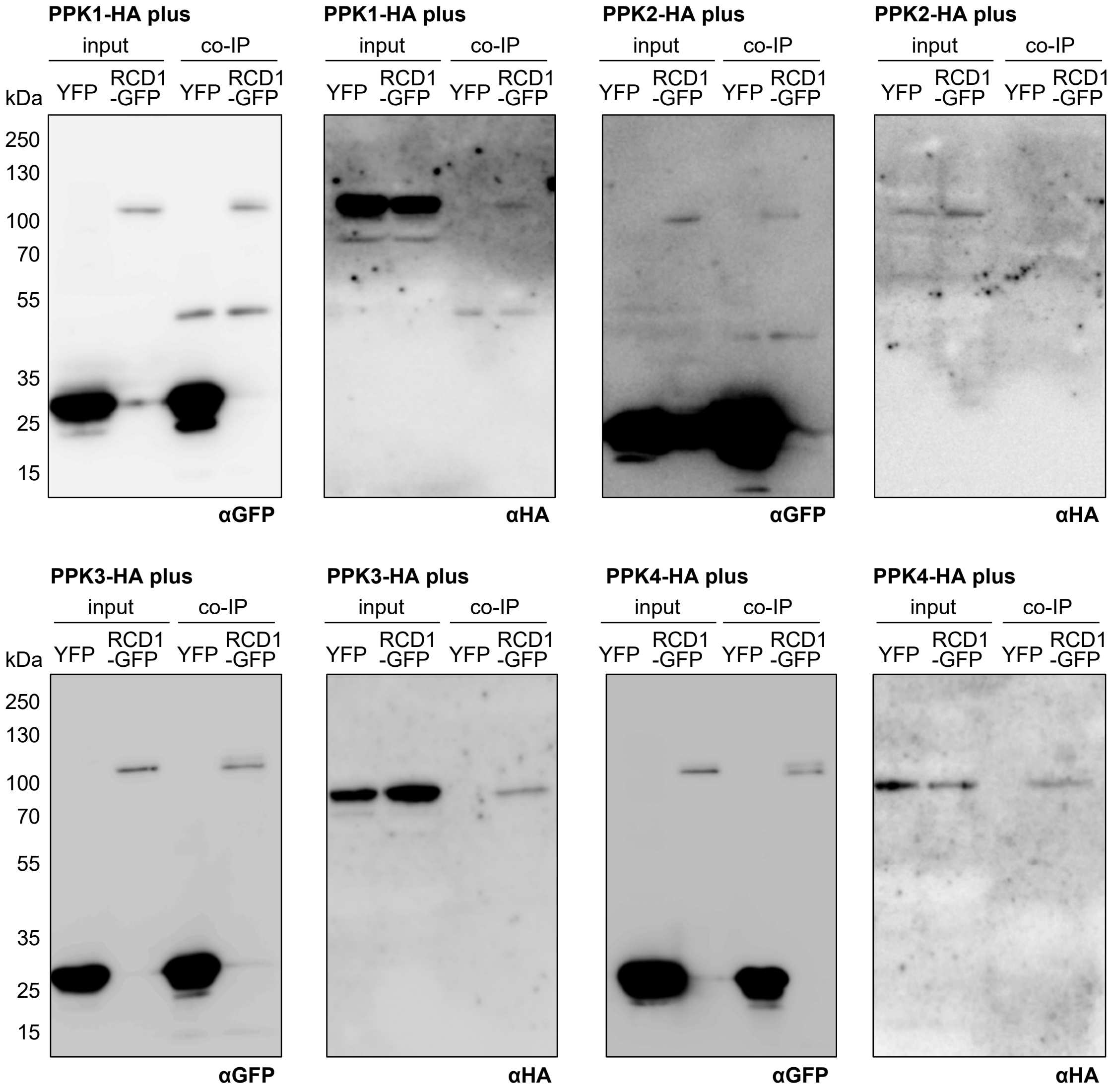


**Supplementary figure 5. RCD1-GFP interacts with PPK-HA in tobacco.**

RCD1-GFP was transiently co-expressed with HA-tagged versions of PPK1, 2, 3 or 4 in *N. benthamiana*. YFP served as negative control. At 72 hours post infiltration, RCD1-GFP and YFP were immunoprecipitated with GFP-specific antibody and co-precipitating PPK-HA proteins were detected by αHA immunoblot. Immunoprecipitation of RCD1-GFP and YFP was confirmed by an αGFP immunoblot. 'Input' samples were taken before immunoprecipitation and included on the immunoblots to test for equal expression and loading.

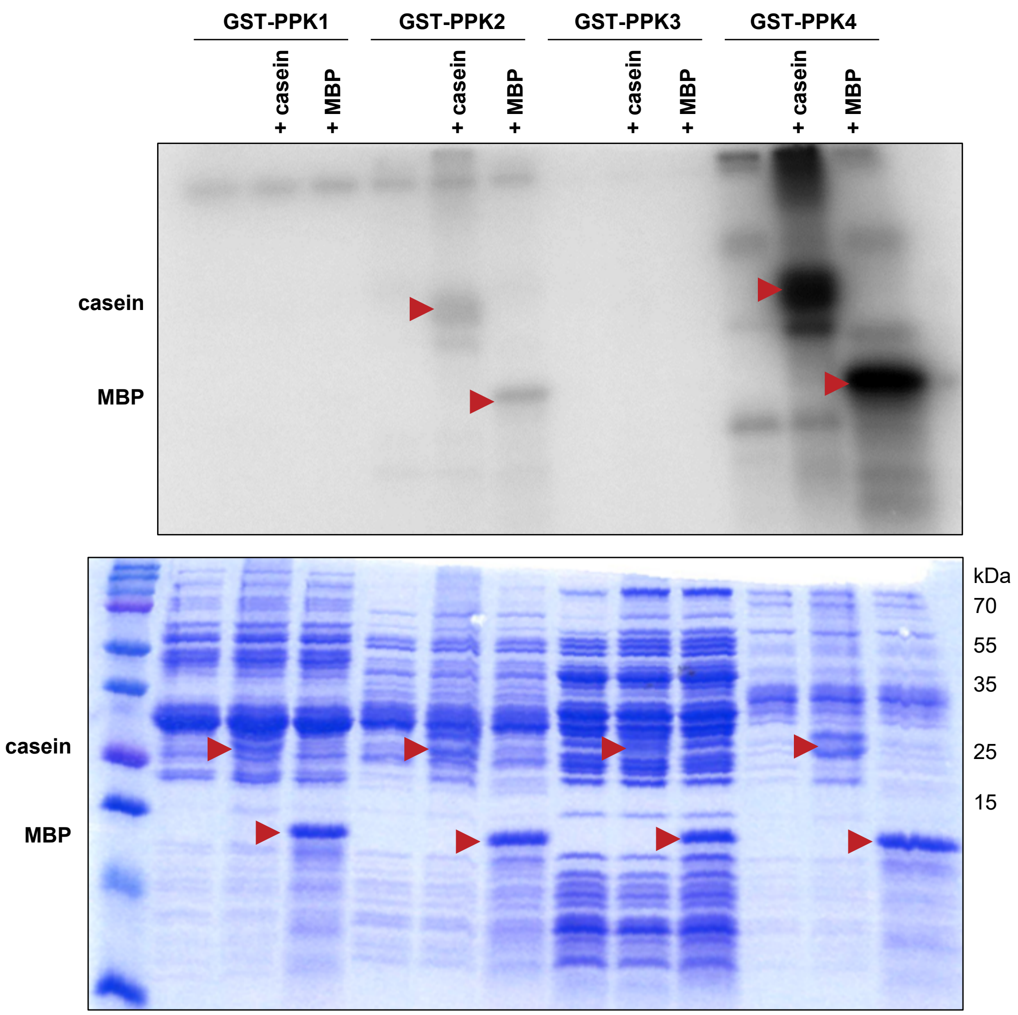


**Supplementary figure 6. Recombinant PPK2 and PPK4 are active in *in vitro* kinase assays.**
Recombinant GST-PPK1-4 were used together with generic substrates casein and myelin basic protein (MBP) in an *in vitro* kinase assay. Upper panel shows autoradiograph, lower panel shows the Coomassie-stained SDS-PAGE.

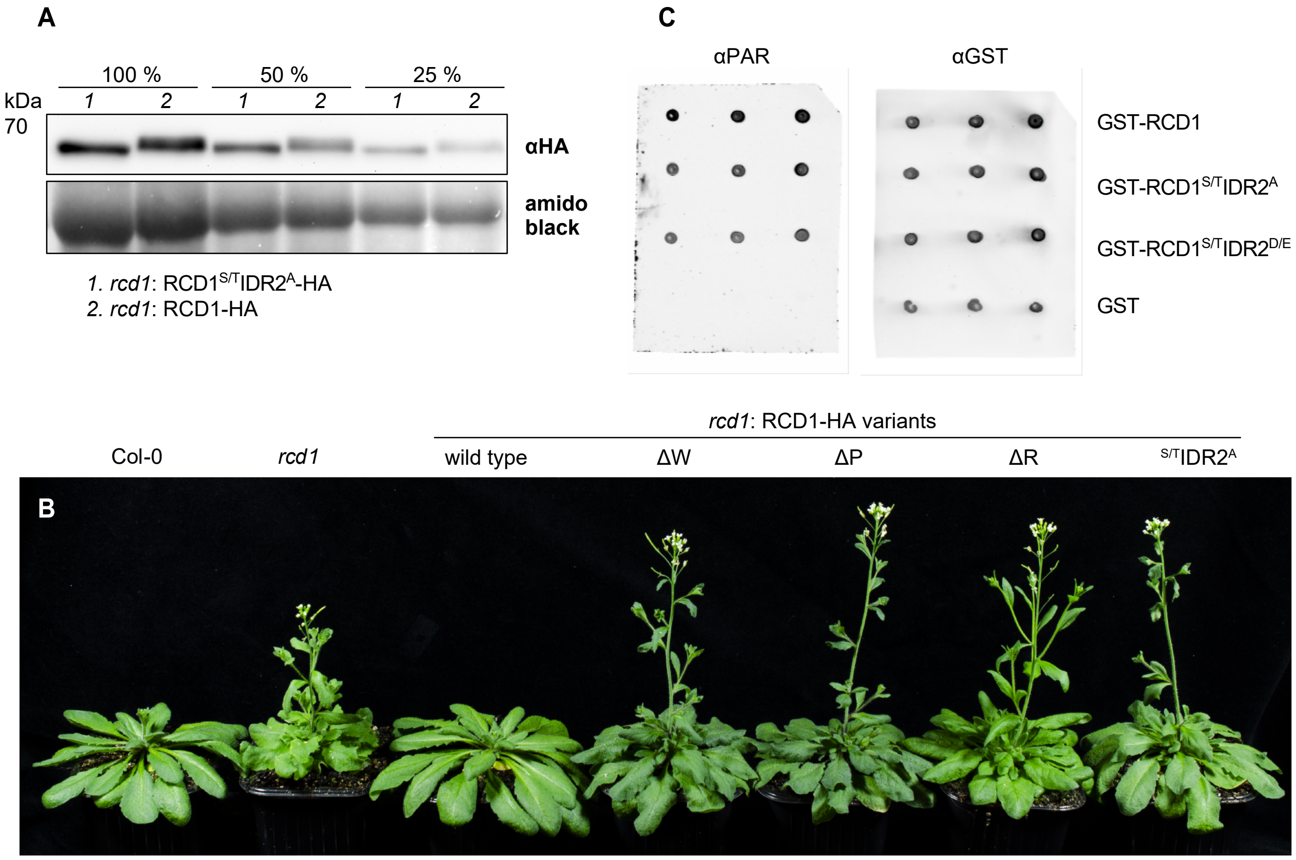


**Supplementary figure 7. Effect of RCD1 phosphorylation in IDR2 by PPKs on the protein function.**
**A.** *In vivo* phosphorylation pattern of RCD1$^{S/T}$IDR2$^{A}$-HA is different from that of the wild type RCD1-HA. Upper band corresponding to phosphorylated form of RCD1 is not detectable in RCD1$^{S/T}$IDR2$^{A}$-HA line as visualized by immunoblot analysis of protein extracts with HA-specific antibody. The lines with approximately equal expression of RCD1 were selected for this comparison. 100% corresponds to 100 µg of total protein. Rubisco large subunit detected by amido black staining is shown as a control for equal protein loading.
**B.** Early flowering phenotype of *rcd1* is not restored by expression of RCD1$^{S/T}$IDR2$^{A}$-HA in *rcd1* background. Picture shows 5-week-old plants of representative lines under standard growth conditions. Domain deletion lines are shown in the same figure for comparison.
**C.** PPK phosphorylation in IDR2 does not affect PAR binding by RCD1 *in vitro*. PAR binding activity of immobilized GST-tagged RCD1$^{S/T}$IDR2$^{A}$ and RCD1$^{S/T}$IDR2$^{D/E}$, mimicking non-phosphorylated or fully phosphorylated RCD1, respectively, was accessed by dot-blot assay using PAR-specific antibody. GST was used as a negative control. GST antibody was used to assess protein loading.

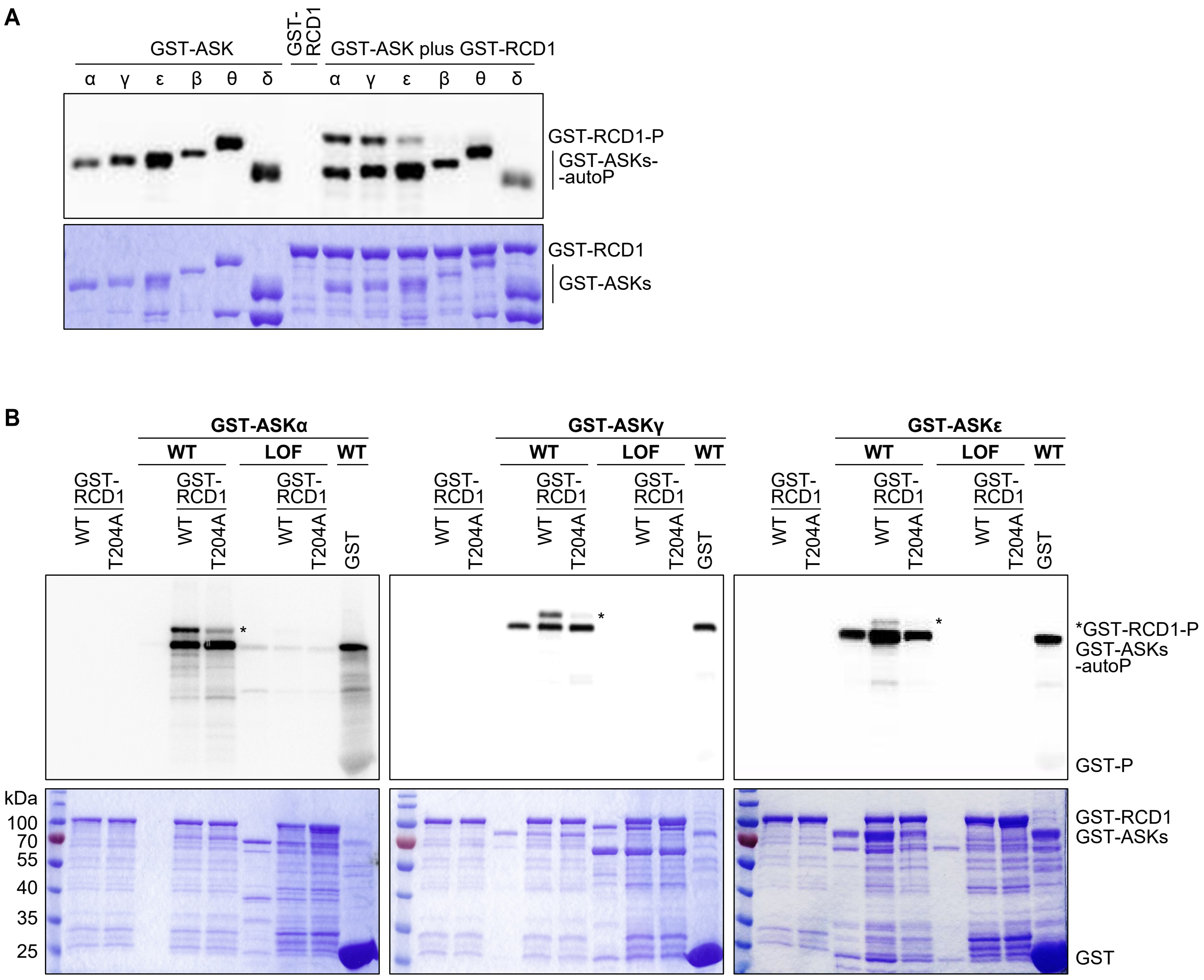


**Supplementary figure 8. ASKα, ASKγ, and ASKε phosphorylate RCD1 *in vitro*.**

**A.** Specificity of ASKα, ASKγ and ASKε towards RCD1. Recombinant ASK-GSTs were used together with recombinant GST-RCD1 protein in an *in vitro* kinase assay. P – phosphorylated protein; autoP – autophosphorylated protein.

**B.** Thr204 is the target for ASKs. ASKα,γ,ε-GST were used with recombinant GST-RCD1 or GST-RCD1T204A in an *in vitro* kinase assay. LOF indicates loss-of-function constructs of ASKs. Upper panels show autoradiographs, lower panels show the Coomassie-stained SDS-PAGE.
